# MViewEMA: Efficient Global Accuracy Estimation for Protein Complex Structural Models Using Multi-View Representation Learning

**DOI:** 10.1101/2025.07.25.666906

**Authors:** Dong Liu, Xuanfeng Zhao, Tianyou Zhang, Lei Xie, Enjia Ye, Fang Liang, Haodong Wang, Guijun Zhang

**Affiliations:** College of Information Engineering, Zhejiang University of Technology, 288 Liuhe Road, 310023 HangZhou, China

## Abstract

Estimation of model accuracy (EMA) is crucial for the reliable application of protein structure models in biological research. With the rapid advancement of protein structure prediction techniques and the explosive growth of predicted structural data, existing EMA methods struggle to balance computational efficiency with estimation performance. Here, we present MViewEMA, a single-model EMA method that leverages a multi-view representation learning framework to integrate residue-residue interaction features from micro-environment, meso-environment, and macro-environment levels for global accuracy assessment of protein complex models. Benchmark results demonstrate that MViewEMA outperforms state-of-the-art EMA methods in global accuracy assessment, achieving more than a 10-fold improvement in computational efficiency compared to our previous method, DeepUMQA3. This method enables efficient selection of high-quality protein complex models from large-scale structural datasets and achieved top performance in model selection tracks during the CASP16 blind test, demonstrating its potential to enhance the accuracy of complex structure prediction when integrated into modern frameworks such as AlphaFold-Multimer, AlphaFold3, and DiffDock-PP.

## INTRODUCTION

The emergence of protein structure databases with hundreds of millions of predicted models (e.g., AlphaFold DB^1^ and ESM Metagenomic Atlas^2^) has driven rapid progress in structural biology and bioinformatics^3^. These large-scale databases pose a challenge for accurately assessing structural quality to ensure their reliable application in life science research. This challenge becomes particularly acute in the post-AlphaFold era, where structure prediction may evolve into a model selection problem, requiring efficient discrimination of high-accuracy structures from extensive conformational ensembles^4^. The recent introduction of a dedicated massive model selection track^4^ in CASP16 exemplifies the growing demand for efficient estimation of model accuracy (EMA)^5, 6^ methods to facilitate the practical application of these structural resources.

EMA provides quantification of the deviation between predicted protein models and their native structures when experimental structural data are not yet available, establishing accuracy metrics for model selection^7^. With the advancement of structure prediction methodologies, EMA techniques have increasingly become intrinsic self-assessment modules integrated into modern prediction frameworks^8–10^. These frameworks leverage intermediate representations or prior knowledge derived from the modeling process to evaluate structure accuracy, as exemplified by sequence-based methods such as AlphaFold^8, 9^ and structure-based docking protocols^11–14^. For sequence-based methods, AlphaFold2^8^ derives its self-assessment scores from the single and pair representations of multiple sequence alignments (MSAs) and structural templates, while AlphaFold3^9^ employs an enhanced four-block confidence module that utilizes corresponding representations to evaluate and select diffusion-generated models. For structure-based protocols, diffusion-driven docking methods such as DiffDock-PP^11^ and template/energy-based methods like HDOCK^12^ utilize modeling-process information as input for assessing docking model accuracy. While these self-assessment modules demonstrate strong performance within their modeling frameworks, their reliance on framework-specific input features often limit generalizability across diverse prediction methods.

To address the limitations of self-assessment modules, third-party EMA approaches (e.g. consensus, quasi-single, and single-model techniques) remain essential for protein structure prediction. Consensus methods^15–18^ evaluate model accuracy by identifying common structural features across multiple models, such as MULTICOM^19^, making their performance dependent on the quality of the input model pool. MULTICOM^19^ has been one of the pioneering forces in the field, consistently excelling in CASP experiments and advancing the development of consensus-based quality assessment strategies. To mitigate the reliance on quality of input model pool, quasi-single-model methods^20–22^ employ internal structure prediction techniques to generate reference structures and compare them with candidate models using deep learning or structural alignment. ModFOLD^22^ is a representative method in this category, maintaining state-of-the-art (SOTA) performance while providing freely accessible EMA services to a wide range of bioscience communities. Single-model methods^23–27^ provide a fully autonomous solution for evaluating protein model accuracy, requiring only a single model as input. By employing either statistical potential-based scoring systems or machine learning frameworks, these approaches eliminate the need for input model ensembles or external reference structures. Among them, ProQ^28^ is one of the first methods to apply machine learning techniques to protein EMA, laying the foundation for the development of single-model approaches.

Single-model methods represent the foundational paradigm of EMA. These approaches have established field dominance through three defining characteristics: First, their methodology-independent nature eliminates any reliance on specific modeling algorithms or reference models, enabling universal evaluation of structures generated by techniques ranging from traditional docking-based protocols (e.g., HDOCK) to end-to-end deep learning frameworks (e.g., AlphaFold series). Second, their inherent computational efficiency, requiring only single model input, provides distinct advantages for large-scale model selection across diverse prediction methods. Most critically, their intrinsic evaluation paradigm performs direct structural feature analysis, achieving EMA’s primary objective of property-based assessment, while maintaining rigorous theoretical standards. These core advantages have cemented single-model methods as the central driver of EMA advancement, where evolution from statistical potentials to deep learning frameworks constitutes a continuous refinement of this paradigm, solidifying its position as the gold standard for protein EMA.

In recent years, single-model methods have made significant progress driven by advancements in deep learning technologies, such as DeepAccNet-MSA^29^, DeepUMQA3^30^ and VoroIF-GNN^31^. DeepAccNet-MSA^29^ leverages co-evolutionary information derived from MSA and convolutional architectures to predict the local accuracy of monomer models, making it the top-performing single-model local EMA method in CASP14. With the advancement of protein language models, our developed method DeepUMQA3^30^ further incorporates evolutionary features extracted by ESM (i.e., Evolutionary Scale Modeling)^32^, along with intra-chain and inter-chain USR^25^ descriptors, to assess the local accuracy of complex models, achieving SOTA performance in interface-level local EMA in CASP15. In contrast to these evolution-aware methods, VoroIF-GNN^31^ adopts a purely structure-based strategy without relying on any evolutionary information. It derives interface contacts from the Voronoi tessellation of atomic spheres to construct a contact graph and predicts the accuracy of each contact using an attention-based graph neural network, demonstrating top performance for multimeric model selection in CASP15.

Most high-performing single-model methods rely on evolution-aware features derived from MSAs, structural templates, or protein language models. Since these features are commonly used for structure prediction, such methods may exhibit bias towards protein structures that share similar evolutionary information, especially for targets with weak evolutionary signals (e.g., antibody–antigen). Extracting these features is computationally intensive and limits the applicability of these methods in large-scale, high-throughput model evaluation. While some single-model methods have attempted to assess model accuracy using purely structure-based features, they have generally struggled to achieve SOTA performance. In addition, many existing EMA methods focus on residue-level or interface-level accuracy, making them less effective in assessing the global structural integrity from comprehensive perspectives, which is critical for capturing the overall correctness of domain-domain or inter-chain arrangements^8^ and model ranking^9^. Thereby, developing an efficient, accurate, and generalizable single-model EMA method that is independent of the structure modelling process remains a critical challenge in structural bioinformatics.

To address these limitations, building upon our previous work (DeepUMQA series)^25, 30^, we further developed a single-model EMA framework based on multi-view representation learning^33^, MViewEMA, enabling efficient global confidence estimation and model selection in large-scale structural sampling scenarios. The proposed method extracts residue–residue interaction features from three complementary views, including micro-environment, meso-environment, and macro-environment. Each feature view is processed by dedicated networks using graph attention^34^, convolutional^35^, and Transformer^36^ architecture, whose outputs are integrated through cross-view information flow to predict the global confidence score. MViewEMA operates without reliance on modeling-driven information sources (e.g., MSAs, templates or protein language models), establishing analytical independence from the modeling process. This enables a fully decoupled accuracy assessment solution for a wide range of protein structure prediction methods.

Benchmark results demonstrate that MViewEMA achieves strong accuracy and efficiency compared to SOTA single-model methods on both the CASP15 benchmark dataset and the CASP16 EMA blind test, with a 10.24-fold increase in runtime efficiency over our previously developed DeepUMQA3.

## RESULTS

### MViewEMA Overview

MViewEMA is a deep learning-based computational protocol developed for global accuracy estimation of protein models. Given a protein complex model, it employs a multi-view representation learning framework to comprehensively capture the relationship between model accuracy and protein features across micro to macro scales. The framework integrates three views of residue – residue interaction features, including <u>mi</u>cro-<u>e</u>nvironment (MiE), <u>me</u>so-<u>e</u>nvironment (MeE), and <u>ma</u>cro-<u>e</u>nvironment (MaE), which are processed by specialized heterogeneous network architectures comprising graph, convolutional, and transformer modules to predict a global confidence score (i.e., TM-score^37^) of the entire structure (**Fig. 1**). Specifically, (1) The MiE features encode amino acid properties (e.g., residue type encoding and physicochemical properties), geometric configurations (e.g., torsion angle, secondary structure, backbone bond angle and length), and statistical features (e.g., one-body energy terms) of individual residues, providing fine-grained protein descriptions. These MiE features are further enriched by incorporating residue substitution probabilities^38^ from the MeE features and inter-residue distance map from the MaE features, and are processed using graph attention networks (GATs)^34^. Through iterative message passing between residues, the GATs progressively aggregate features from individual residues to the entire protein, improving expressive capacity from micro-level descriptors to macro-level structure awareness. (2) The MeE features simulate the local atomic environment surrounding each residue. A voxelized 3D grid centered on each residue is constructed, and cascaded 3D CNNs^35^ are employed to extract local spatial features. This enables the method to capture neighborhood-level spatial dependencies and serves as a transitional bridge toward global structural understanding. (3) The MaE features capture the geometric relationships and statistical dependencies between each residue and all other residues in the protein, including two-body energy terms, residue frame alignment encoding, distance and orientation maps. These MaE features with MeE-2D representations are processed by 2D convolutions and pooling layers, followed by biaxial attention-based Transformer networks^36^. This design enables effective learning of long-range residue– residue dependencies that are crucial for improving overall EMA accuracy. Finally, the representations from MiE, MeE, and MaE are integrated via matrix multiplication and pooling layers, enabling MViewEMA to jointly infer across multiple structural scales and output a global confidence score for the protein model. Full details for both multi-view features and network architecture are provided in Methods.

**Fig. 1.**
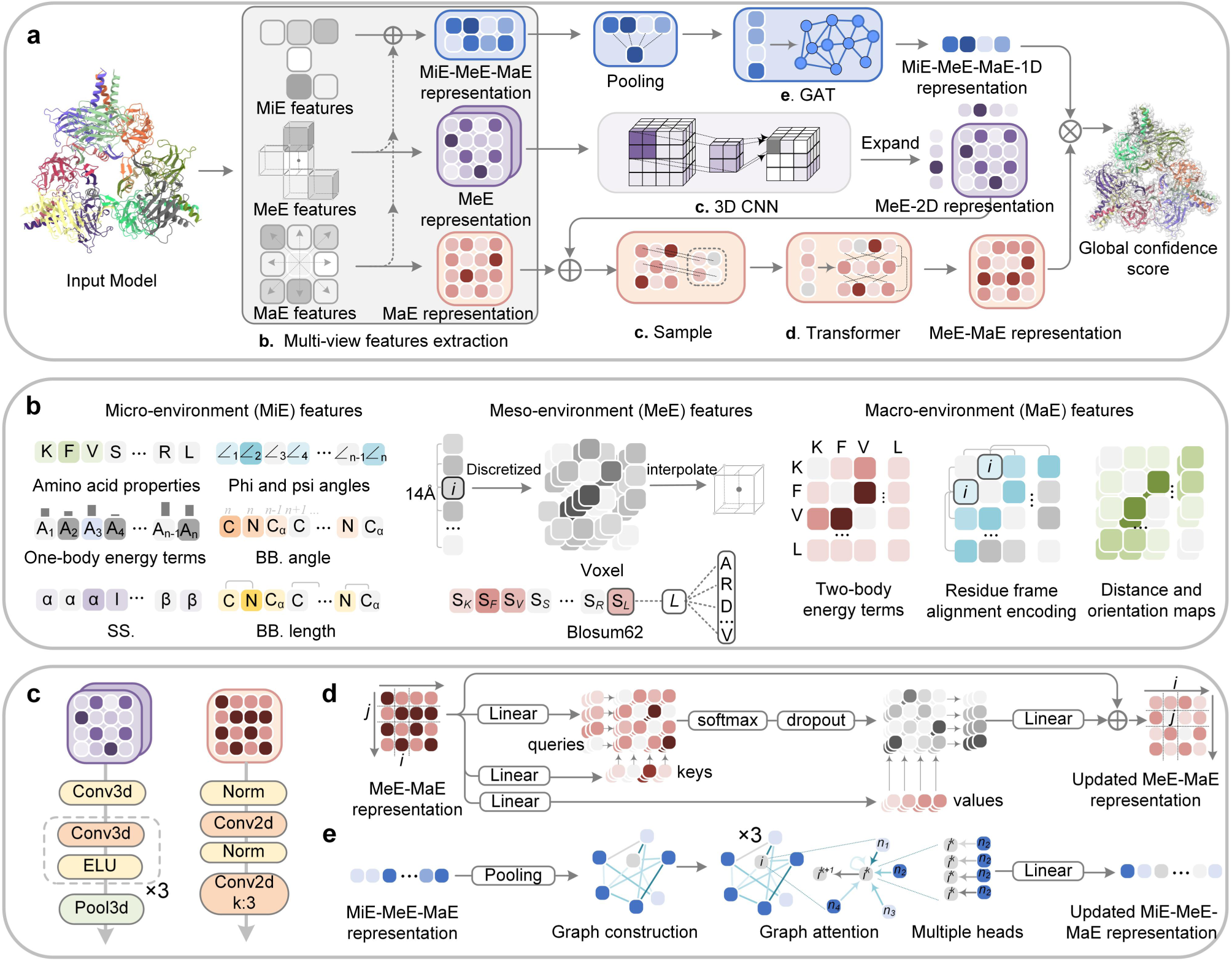
Overview of the MViewEMA framework for global accuracy assessment of protein models. **a**, Overall architecture of MViewEMA. Given an input protein model, multi-view residue–residue interaction features are extracted, including micro-environment (MiE), meso-environment (MeE), and macro-environment (MaE). These features are hierarchically aggregated through a combination of convolution, transformer, and graph modules to predict a global confidence score. **b**, Illustration of the three residue–residue interaction features. (1) MiE features encode the physicochemical, geometric, and statistical attributes of individual residues; (2) MeE features represent the geometric context of each residue within its local atomic neighborhood and incorporate residue substitution probabilities; (3) MaE features describe the distance, spatial orientation, and statistical energy relationships between each residue and all other residues within the protein. **c**, The MeE representation is passed through cascaded 3D CNN layers to extract local spatial features (left panel). The MaE and MeE-2D representations are processed by 2D convolution and pooling layers for spatial downsampling (right panel). **d**, The MeE-MaE representation is processed by a transformer network with a biaxial multi-head attention mechanism. **e**, The MeE-MiE-MaE representation is fed into multi-head graph attention networks which update the representation through iterative message passing.

### Comparison between MViewEMA and state-of-the-art methods on the benchmark dataset

To test the global accuracy assessment performance of the proposed method, MViewEMA is systematically compared with all 16 single-model methods and top 5 consensus methods of CASP15 on the official dataset. The dataset and the training dataset of MViewEMA were strictly separated in time, and a 40% sequence identity cutoff was applied at the complex-chain level in the training data to prevent potential information leakage (see training and test datasets in Methods). To ensure a rigorous performance comparison, while MViewEMA was trained using TM-score as the global confidence score, we followed CASP protocols to analyze global EMA performance based on both TM-score and Oligo-GDTTS^39^. All data were obtained from official CASP repository, and the analysis was conducted using publicly available code provided by the CASP organizers (https://git.scicore.unibas.ch/schwede/casp15_ema), with an assessor-defined sum Z-score formula that integrates Pearson correlation, Spearman correlation, ROC AUC, and Top1Loss^6^ (**Supplementary Note S1**).

MViewEMA shows top performance in global accuracy estimation on the CASP15 benchmark dataset (**Fig. 2**). It outperformed all 16 single-model methods, achieving the Z-score^39^ (TM-score) of 4.11 and Z-score (Oligo-GDTTS) of 3.67, significantly surpassing the second-best method, Manifold, which achieved Z-scores of 3.03 and 3.39, respectively (**Fig. 2a**). Compared to our previously proposed methods, GraphGPSM^26^ (Group name: GuijunLab-Threader) and DeepUMQA3^30^ (Group name: GuijunLab-RocketX), MViewEMA improved the sum Z-score on TM-score and Oligo-GDTTS by at least 1.91 and 1.94, respectively. This performance gain is primarily attributed to the introduction of multi-view representation learning framework in MViewEMA, which maps protein structure to model accuracy from different views, including MiE, MeE, and MaE. Furthermore, we compare the performance of MViewEMA and the top 5 single-model methods in terms of model selection (Top1Loss) and accuracy discrimination (ROC AUC), as shown in **Fig. 2b and Supplementary Fig. S1 and Tables S1-S2**. For model selection, MViewEMA achieved the best Top1Loss on both TM-score and Oligo-GDTTS (0.104 and 0.236), significantly outperforming Manifold (0.144 and 0.295) and GraphGPSM (0.187 and 0.399). Consistent trends were observed in model accuracy identification, where MViewEMA also achieved the highest average ROC AUC for these metrics (0.704 and 0.688), as shown in **Fig. 2b**. However, MViewEMA did not achieve the best results in Pearson and Spearman correlations (**Supplementary Tables S1-S2**). Further analysis found that MViewEMA tended to overestimate global confidence scores, likely due to the inclusion of monomer structure data during training and a distribution bias toward high-quality models in the dataset. Nevertheless, incorporating monomer data enables the network to learn fundamental folding patterns and physicochemical properties of individual chains, enhancing its generalizability and assessment accuracy for complex models, as supported by our ablation studies (**Fig. 6**).

**Fig. 2.**
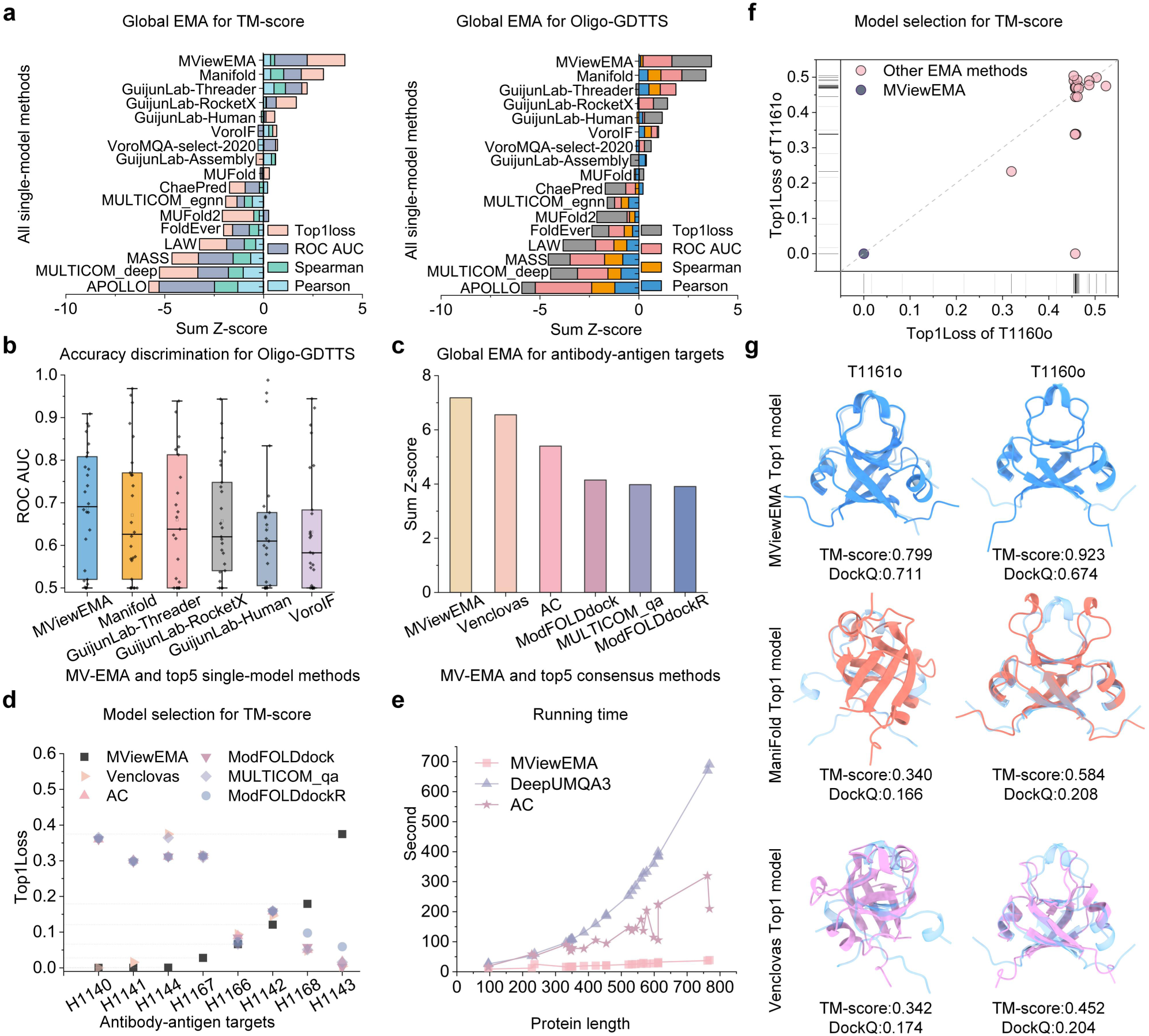
Performance comparison between MViewEMA and SOTA EMA methods on benchmark datasets. **a**, Global accuracy assessment of MViewEMA and all 16 single-model methods for TM-score (left panel) and Oligo-GDTTS (right panel) metrics on the CASP15 dataset. Sum Z-score integrates Top1Loss, ROC AUC, Spearman, and Pearson correlation. **b**, Comparison of ROC AUC across CASP15 targets for MViewEMA and the top 5 single-model methods. **c**, Sum Z-score of MViewEMA and the top 5 consensus methods on antibody–antigen targets. The Z-score, defined as in **a**, combines TM-score and Oligo-GDTTS. **d**, Per-target Top1Loss of MViewEMA and top 5 consensus methods for antibody–antigen targets. **e**, Running time of single-model methods (MViewEMA and DeepUMQA3) and the consensus method AssemblyConsensus (AC). **f**, Top1Loss of MViewEMA and all other EMA methods on the challenging dimers T1160o and T1161o. **g**, Structures of top-ranked models selected by MViewEMA, ManiFold, and Venclovas for T1160o and T1161o. The native structure is shown in light blue, while the predicted structures are depicted in distinct colors.

To further evaluate its performance in challenging scenarios with high structural diversity and scarce homologous templates, MViewEMA was compared against the top 5 consensus methods on antibody – antigen complex models (**Figs. 2c-d**). It achieved the highest sum Z-score of 7.183 in global accuracy assessment, outperforming consensus methods and surpassing the best-performing method MULTICOM_qa^16^ by 3.98. As a single-model method, MViewEMA is independent of model pool quality and effectively compensates for the limitations of consensus methods. While MULTICOM_qa performed excellently in all CASP15 targets, its performance on antibody– antigen targets was suboptimal, primarily due to the limitation of most structure prediction methods for weakly homologous targets, which resulted in low-quality model pools (e.g., only 3.4% (11/319) for antibody–antigen target H1140 had TM-score > 0.7). MViewEMA selected top1 models that were close to or matched the best accuracy in 6 antibody–antigen targets and achieved the best average Top1Loss (0.096) across all antibody–antigen targets (**Fig. 2d**). It is a 23.2% improvement over the second-best method Venclovas^40^ (Top1Loss = 0.125), which demonstrated top performance in CASP15 complex structure prediction (ranked second). These findings suggest that MViewEMA exhibits broader applicability to antibody– antigen models than consensus methods, and may offer practical advantages for downstream applications such as antibody design.

Given the challenges associated with selecting high-quality models from structural pools containing mostly low-quality protein models, we analyzed the performance of all EMA methods on two challenging homomers, T1160o and T1161o, which are difficult for both structure prediction and accuracy estimation (**Fig. 2f**). For these targets, the average TM-score of the model pools were 0.334 and 0.469, respectively. Except for MViewEMA, no other EMA method successfully identified the best model for both targets, typically selecting low-quality models instead. Notably, only our previously developed DeepUMQA3 successfully selected the best model for one target T1161o. Based on DeepUMQA3, the improved version MViewEMA selected the best models for both targets, further validating the effectiveness of introducing multi-view representation learning. **Fig. 2g** shows the top1 model selected by different top-performance EMA methods, where the model selected by MViewEMA outperformed those selected by the single-model method ManiFold and consensus method Venclovas in both TM-score and DockQ^41^.

To verify the efficiency of MViewEMA, we compared its runtime performance with that of the single-model method DeepUMQA3 and the consensus method AssemblyConsensus (AC). Under the same hardware configuration (Intel 2.10 GHz, single-core CPU), MViewEMA achieves an average runtime of 22.63 seconds for each protein model, compared to 235.96 seconds for DeepUMQA3 and 120.10 seconds for AC, corresponding to speedups of 10.43-fold and 5.31-fold, respectively, as shown in **Fig. 2e**. Moreover, the runtime of MViewEMA remained relatively stable across a wide range of protein lengths, underscoring its robustness and scalability for handling protein of varying sizes. These above results demonstrate that MViewEMA not only offers strong estimation performance but also excels in efficiency, making it well suited for large-scale protein model evaluation in practical applications.

### Comparison between MViewEMA and AlphaFold self-assessment

To test the potential of MViewEMA to improve structure prediction accuracy in AlphaFold pipelines, we constructed AlphaFold benchmark datasets based on CASP15 targets (see the test datasets in Methods), followed by a comparative analysis against the self-assessment results of AlphaFold-Multimer^42^ and AlphaFold3^9^. As the AlphaFold framework primarily relies on a TM-score-based confidence metric (i.e., ranking_score) to rank predicted models, and this study focuses on assessing global structural accuracy, we compared MViewEMA with confidence score in terms of both accuracy estimation and model selection.

On the datasets of AlphaFold-Multimer and AlphaFold3 with default top 5 models, MViewEMA performs comparably to self-assessment of AlphaFold-Multimer and AlphaFold3 in terms of protein model selection (**Supplementary Table S3**). The average TM-score of the top-ranked models selected by MViewEMA were 0.777 and 0.773, closely matching those selected by AlphaFold-Multimer (0.772) and AlphaFold3 (0.771). Notably, the average best structures observed in the model pools were 0.787 (AlphaFold-Multimer) and 0.802 (AlphaFold3), indicating that while AlphaFold3 was able to achieve higher prediction accuracy, its self-assessment failed to reliably identify them. These results highlight the importance of not only improving the structure prediction itself but also enhancing the performance of model accuracy estimation. Since both datasets contain only five default models for each target, the limited number of candidate structures leads to relatively minor differences in model selection performance. Therefore, we conducted further analyses on datasets with larger model pools, AFM Dataset-MT (for AlphaFold-Multimer) and AF3 Dataset-Seeds (for AlphaFold3), to better test the effectiveness of MViewEMA.

For AlphaFold-Multimer, MViewEMA demonstrates significantly improved performance over its self-assessment in terms of global accuracy assessment on AFM Dataset-MT, as illustrated in **Figs. 3a and 3d**. On the test set, MViewEMA achieves a Top1Loss of 0.095 in TM-score, which is better than AlphaFold-Multimer’s 0.113. Among all targets, MViewEMA outperforms AlphaFold-Multimer in 21 out of 26 targets in Top1Loss (**Fig.3d (1)**). The median TM-score of the top1 models selected by MViewEMA reaches 0.886, representing an improvement of about 12% over those selected by AlphaFold-Multimer self-assessment (median TM-score = 0.791), as shown in **Supplementary Fig. S2**. For antibody–antigen targets, the average accuracy of the top1 models selected by MViewEMA is TM-score = 0.814, which is markedly higher than TM-score = 0.740 obtained by AlphaFold-Multimer self-assessment (**Fig. 3b**). This result further supports the notion that for targets with limited evolutionary information, self-assessment of structure predictions relying on modeling-driven features (e.g. MSAs or templates) may struggle to identify high-quality structures, whereas MViewEMA independent of the modeling process can achieve superior model selection performance. A successful result example of MViewEMA from target H1143 is shown in **Fig. 3e**. The top1 structure selected by MViewEMA achieves TM-score = 0.961 and DockQ = 0.774, whereas AlphaFold-Multimer selection only yields TM-score = 0.637 and DockQ = 0.041, with an incorrectly predicted interaction interface. Additionally, in terms of other analytical metrics, MViewEMA also performed well, achieving ROC AUC = 0.686, Pearson correlation = 0.456, and Spearman correlation = 0.430, all of which are significantly higher than the corresponding values for AlphaFold-Multimer self-assessment (0.629, 0.352, and 0.335), as shown in **Figs. 3a and 3d (2-4)**. Particularly, MViewEMA achieves relative improvements of 29.5% and 28.4% in Pearson and Spearman correlations, respectively.

**Fig. 3.**
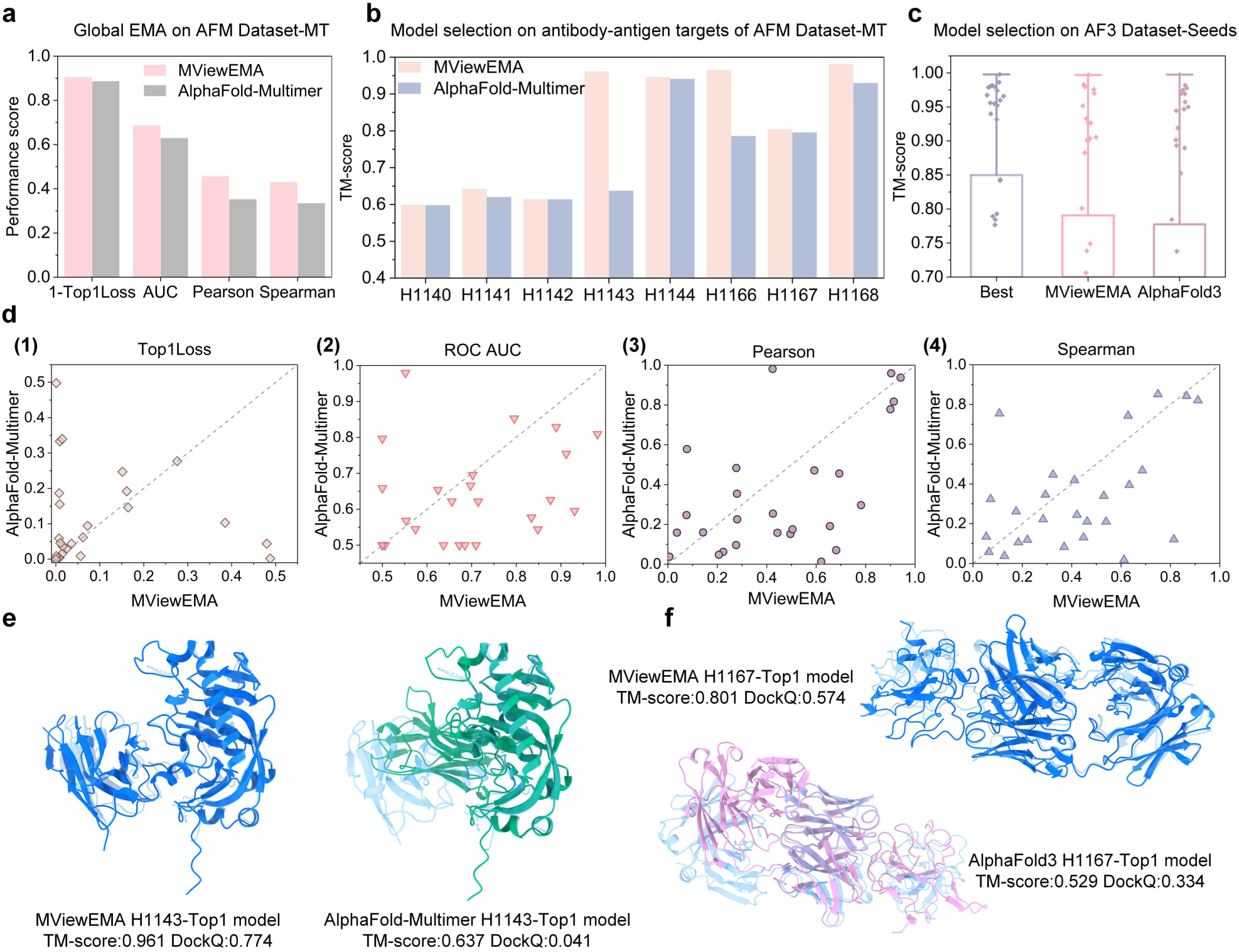
Performance comparison of MViewEMA with AlphaFold self-assessment. **a**, Global accuracy assessment performance of MViewEMA and AlphaFold-Multimer on AFM Dataset-MT. **b**, TM-score of top1 models selected by MViewEMA and AlphaFold-Multimer for each antibody-antigen target. **c**, TM-score of best available models (Best), MViewEMA-selected models, and AlphaFold3-selected models on AF3 Dataset-Seeds. **d**, Performance comparison between MViewEMA and the self-assessment of AlphaFold-Multimer across four analytical metrics: Top1Loss **(1)**, ROC AUC **(2)**, Pearson correlation **(3)**, and Spearman correlation **(4)**. **e**, Structures of top-ranked models selected by MViewEMA and AlphaFold-Multimer for the antibody–antigen target H1143. The native structure is shown in light blue, while the selected structures are depicted in distinct colors. **f**, Structures of top-ranked models selected by MViewEMA and AlphaFold3 for the antibody–antigen target H1167. The color scheme is like **e.**

For AlphaFold3, MViewEMA achieves performance comparable to its self-assessment on several global accuracy metrics and outperforms it in model selection on the AF3 Dataset-Seeds, as shown in **Fig. 3c and Supplementary Fig. S2**. On the test set, MViewEMA achieved a better Top1Loss (0.059) than AlphaFold3 self-assessment (0.072), while maintaining a largely consistent distribution of top1 model accuracies. For antibody-antigen targets, the top1 models selected by MViewEMA and AlphaFold3 also show comparable performance, with average TM-score of 0.741 and 0.749, while exhibiting complementary strengths on specific targets (**Fig. 3f**). On target H1167, the top1 model selected by MViewEMA achieved TM-score=0.801 and DockQ = 0.574, significantly outperforming the structure selected by AlphaFold3 self-assessment (TM-score = 0.529, DockQ = 0.334). Conversely, AlphaFold3 self-assessment performed better on target H1142 (**Supplementary Fig. S3**). Additionally, in terms of other analytical metrics, AlphaFold3 self-assessment achieved higher performance in ROC AUC (0.592), Spearman correlation (0.440), and Pearson correlation (0.344), compared to MViewEMA (0.570, 0.306, and 0.234), as shown in **Supplementary Fig. S2**. It is worth noting that both AlphaFold-Multimer and AlphaFold3 self-assessment leverage MSAs and templates information and can only be used to evaluate their own generated models. In contrast, MViewEMA does not use any structure modeling-driven features, aiming to provide an independent and modeling-decoupled EMA approach applicable across different structure prediction methods. This highlights MViewEMA can serve as a complementary approach to AlphaFold-based self-assessments, particularly in scenarios requiring cross-pipeline accuracy estimation.

On the dataset, we further observed limitations in AlphaFold3 self-assessment regarding model selection. While extensive structural sampling improves the best model accuracy from 0.802 (default setting) to 0.850 in TM-score, it also raises the challenge of selecting the best models. As a result, the predicted accuracy (TM-score = 0.777) of AlphaFold3 using multiple random seeds remains nearly identical to that of the default setting (TM-score = 0.771), while MViewEMA can achieve an average accuracy of TM-score = 0.791. This suggests that relying solely on AlphaFold3 self-assessment may limit its ability to fully exploit the potential of high-quality models generated through extensive sampling. Moreover, it highlights that incorporating EMA methods can enhance AlphaFold3’s performance, as even AlphaFold-Multimer integrated with third-party EMA outperformed AlphaFold3 in the CASP16 results^4^. The above results show that MViewEMA can improve the global structure prediction accuracy of both AlphaFold-Multimer and AlphaFold3, particularly in selecting high-quality structures from the large number of antibody – antigen models generated by AlphaFold-Multimer.

### Comparison between MViewEMA and docking method self-assessment

Protein docking methods are indispensable for modeling protein complexes from known monomer structures, enabling accurate prediction of their assembly and interaction interfaces. To validate the effectiveness of MViewEMA in enhancing accuracy of protein docking predictions, we tested its integration with two representative approaches: DiffDock-PP^11^, a deep learning-based diffusion docking method, and HDOCK^12^, a template/energy-based docking method. On the CASP15 dimer and PDB-2024 targets (see the test datasets in Methods), we compared the model selection performance of MViewEMA with the self-assessment of DiffDock-PP and HDOCK.

For DiffDock-PP, MViewEMA significantly improves its prediction accuracy on the dataset. On CASP15 dimer targets, the top1 structures selected by MViewEMA from DiffDock-PP re-docking models achieve an average TM-score and C-RMSD^11^ of 0.725 and 12.64, with the Top1Loss of 0.06 and 6.86. In contrast, the average accuracy of DiffDock-PP selection is 0.652 (TM-score) and 14.48 (C-RMSD), indicating that MViewEMA improves its prediction accuracy by 11.2% in TM-score and 12.7% in C-RMSD (**Fig. 4a and Supplementary Table S4**). The right panel of **Fig. 4a** and **Supplementary Fig. S4** illustrate the proximity of the models selected by MViewEMA and DiffDock-PP to the optimal models in TM-score and C-RMSD. The results show that MViewEMA achieves a Top1Loss below 0.2 for almost all targets (with only one exception), and for targets such as H1106, T1113o, and T1161o, the Top1Loss is close to zero, demonstrating a notable advantage over DiffDock-PP. In particular, on the H1106 target, MViewEMA successfully selected best model from the DiffDock-PP re-docking models that closely matches the native structure, whereas the self-assessment of DiffDock-PP selected a model with incorrect inter-chain orientation (**Fig. 4e**).

**Fig. 4.**
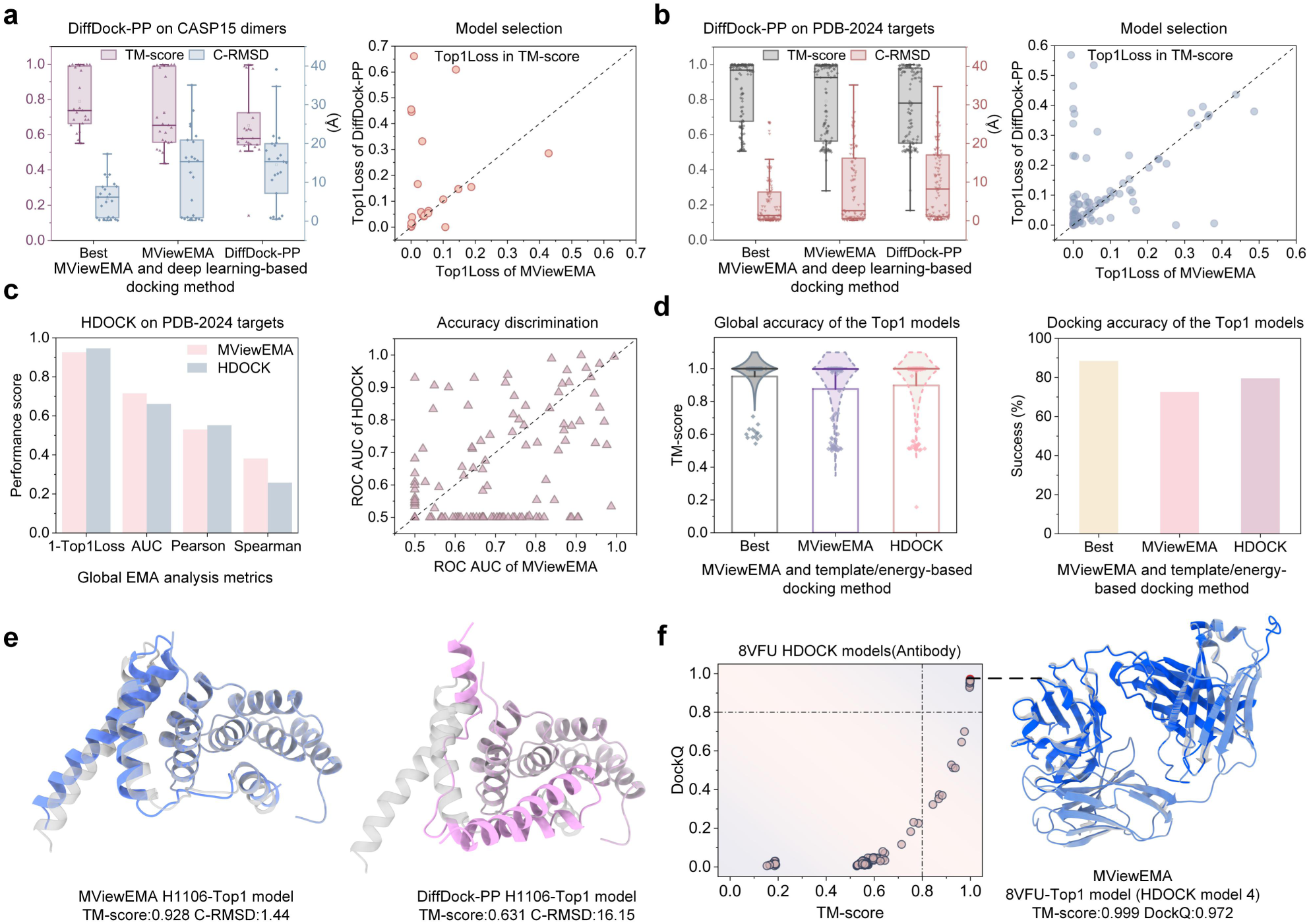
Performance comparison of MViewEMA with DiffDock-PP and HDOCK self-assessment. **a**-**b**, Accuracy (left panel) and loss (right panel) of top1 models selected by MViewEMA and DiffDock-PP self-assessment on CASP15 dimers (**a**) and PDB-2024 targets (**b**). **c**, Global accuracy assessment of analytic metrics (Top1Loss, ROC AUC, Pearson and Spearman correlations) on HDOCK re-docking models of PDB-2024 targets, comparing MViewEMA and HDOCK self-assessment (left panel). The scatter plot illustrates the ROC AUC of both methods for each target (right panel). **d**, TM-score (left panel) and docking success rate (right panel) of top1 models selected by MViewEMA and HDOCK self-assessment on PDB-2024 targets, where docking success is defined as DockQ > 0.23 for protein–protein interfaces. **e**, Structures of top-ranked models selected by MViewEMA and DiffDock-PP self-assessment on target H1106. The native structure is shown in gray, while the selected structures are depicted in distinct colors. **f**, Ture accuracy distribution of 200 HDOCK re-docking models for antibody target 8VFU (left panel) and the structure of top-ranked model selected by MViewEMA (right panel). The color scheme is like **e.**

On DiffDock-PP re-docking models of PDB-2024 targets, the top1 models selected by MViewEMA achieve an average TM-score of 0.791 and C-RMSD of 8.62, outperforming the DiffDock-PP selections (TM-score = 0.766, C-RMSD = 9.59), as shown in **Fig. 4b**, **Supplementary Fig. S5** and **Table S5**. Notably, the median TM-score of the top1 models selected by MViewEMA achieves TM-score = 0.925, which is close to the best model in the dataset (TM-score = 0.968), as shown in the left panel of **Fig. 4b**. Moreover, the top models selected by MViewEMA outperformed the self-assessment results of DiffDock-PP on 67 out of 116 targets (right panel of **Fig. 4b**). While the self-assessment of DiffDock-PP is trained on its own generated models using the protein geometric features and C-RMSD metric, its model selection performance remains less competitive compared to MViewEMA, which was not fine-tuned for DiffDock-PP models. This may be attributed to the fact that MViewEMA not only incorporates geometric features but also integrates physicochemical properties and statistical descriptors, leveraging a multi-view representation framework across MiE, MeE, and MaE to provide a more comprehensive understanding of protein interface characteristics.

For HDOCK, we compared MViewEMA’s performance with its self-assessment score in global accuracy assessment, where the scores (0-0.5 for low quality, 0.5-0.7 for medium quality, 0.7-1 for high quality) are similar to the TM-score metric. It is important to note that the latest version of HDOCK (released on 2023-09-09) postdates the CASP15 targets (2022-12-13), which may potentially lead to data leakage. To ensure the reliability of the test results, our analysis focuses on PDB-2024 complex targets, with CASP15 results provided in **Supplementary Fig. S6**.

On HDOCK re-docking models of PDB-2024 targets, MViewEMA achieves a higher Spearman correlation (0.381) and ROC AUC (0.716) in global accuracy assessment compared to HDOCK self-assessment (Spearman = 0.258, ROC AUC = 0.661), as shown in **Fig. 4c**. Regarding the ROC AUC, MViewEMA and HDOCK were able to distinguish between high-and low-quality models (i.e., ROC AUC > 0.5) on 96 and 69 targets, respectively (right panel of **Fig. 4c**), suggesting that MViewEMA may offer more robust and reliable discrimination. However, MViewEMA performs slightly worse in Pearson correlation (MViewEMA = 0.530 vs. HDOCK = 0.552) and Top1Loss (MViewEMA = 0.074 vs. HDOCK = 0.054). **Fig. 4d** presents the structure prediction performance of MViewEMA and HDOCK in terms of TM-score and docking success rate, where a docking is considered successful if DockQ > 0.23 for protein–protein interfaces. For TM-score, all the top1 models selected by MViewEMA and HDOCK achieve average values of 0.876 and 0.919 (**Supplementary Table S6**), with overall similar distribution of model quality (left panel of **Fig. 4d**). For docking success rate, MViewEMA and HDOCK achieved 72.6% and 79.6% (**Supplementary Table S6**), indicating that HDOCK exhibits an advantage in interface-specific evaluation for its own generated models.

Further analysis of all 116 PDB-2024 targets finds that HDOCK selected top1 models for 90 targets exhibit TM-score > 0.9, reflecting excellent performance on most targets. We speculate that, despite the temporal separation of experimental testing, HDOCK may have implicitly learned docking patterns similar to those in the dataset (analogous to using templates), resulting in an overestimation of structure prediction and accuracy assessment. In contrast, for the 12 medium-quality targets (0.5<TM-score<0.7) of HDOCK, MViewEMA selects top1 models with an average TM-score of 0.547, outperforming HDOCK’s 0.503 (**Supplementary Table S7**). This indicates that MViewEMA is more effective in improving the accuracy of medium-quality model selection compared to HDOCK self-assessment. While MViewEMA shows slightly lower overall model selection performance compared to HDOCK self-assessment, it can still serve as an independent, third-party tool to further verify the reliability of HDOCK selections, as demonstrated in the case of the antibody complex 8VFU (**Fig. 4f**). On this target, only 10% (20 out of 200) of the models are of high quality (TM-score > 0.8), MViewEMA successfully identifies the best model, which is equivalent to the native structure, and its selection is largely consistent with that of HDOCK. Overall, compared to traditional docking methods, MViewEMA may be more suitable for enhancing the structure prediction accuracy of deep learning-based docking methods.

### Comparison of MViewEMA with EMA methods on the CASP16 blind dataset

To more objectively evaluate the performance of EMA methods without modeling-driven information (e.g., MSAs, templates and protein language models), MViewEMA (Group name: Guijunlab-Complex) participated in the community-wide blind test experiment of CASP16 EMA. For a fair comparison on the blind dataset, we focused our analysis on SOTA single-model methods without modeling-driven information (as referenced in the CASP16 ABSTRACT BOOK) on both the global accuracy estimation and model selection tracks. The evaluation involved approximately 14,000 models submitted by structure prediction groups for global accuracy estimation (i.e., QMODE1) and around 500,000 models generated by MassiveFold for model selection (i.e., QMODE3). All test results were obtained from publicly available data provided by the official CASP16 website (https://predictioncenter.org/casp16/index.cgi).

MViewEMA demonstrates superior performance in capturing the correlation between predicted scores and true global model quality, as shown in **Fig. 5a**. MViewEMA achieved the highest average Pearson correlation (Oligo-GDTTS: 0.540, TM-score: 0.472) among all single-model methods without modeling-driven information (**Fig. 5a (1)** and **Supplementary Fig. S7**). Moreover, for all single-model methods, its performance was only surpassed by the best single-model methods G314 and G312, which incorporated MSAs and protein language model features (**Supplementary Fig. S8**). However, for targets with shallow MSAs or low-quality templates, the best single-model methods may struggle to achieve strong performance, such as antibody-antigen targets. For CASP16 antibody-antigen targets, MViewEMA (0.454) performed better than G314 (0.424) and G312 (0.282) in terms of Pearson correlation for TM-score metric (**Supplementary Table S8**). Consistent results were observed in the Spearman correlation (i.e., rank correlation) analysis based on Spearman (**Fig. 5a (2)** and **Supplementary Fig. S8**). Remarkably, even the best-performing single-model method achieved only moderate Spearman values of 0.358 for TM-score and 0.412 for Oligo-GDTTS on all targets. These results indicate that there remains substantial room for improvement in the ranking capability of existing EMA methods.

**Fig. 5.**
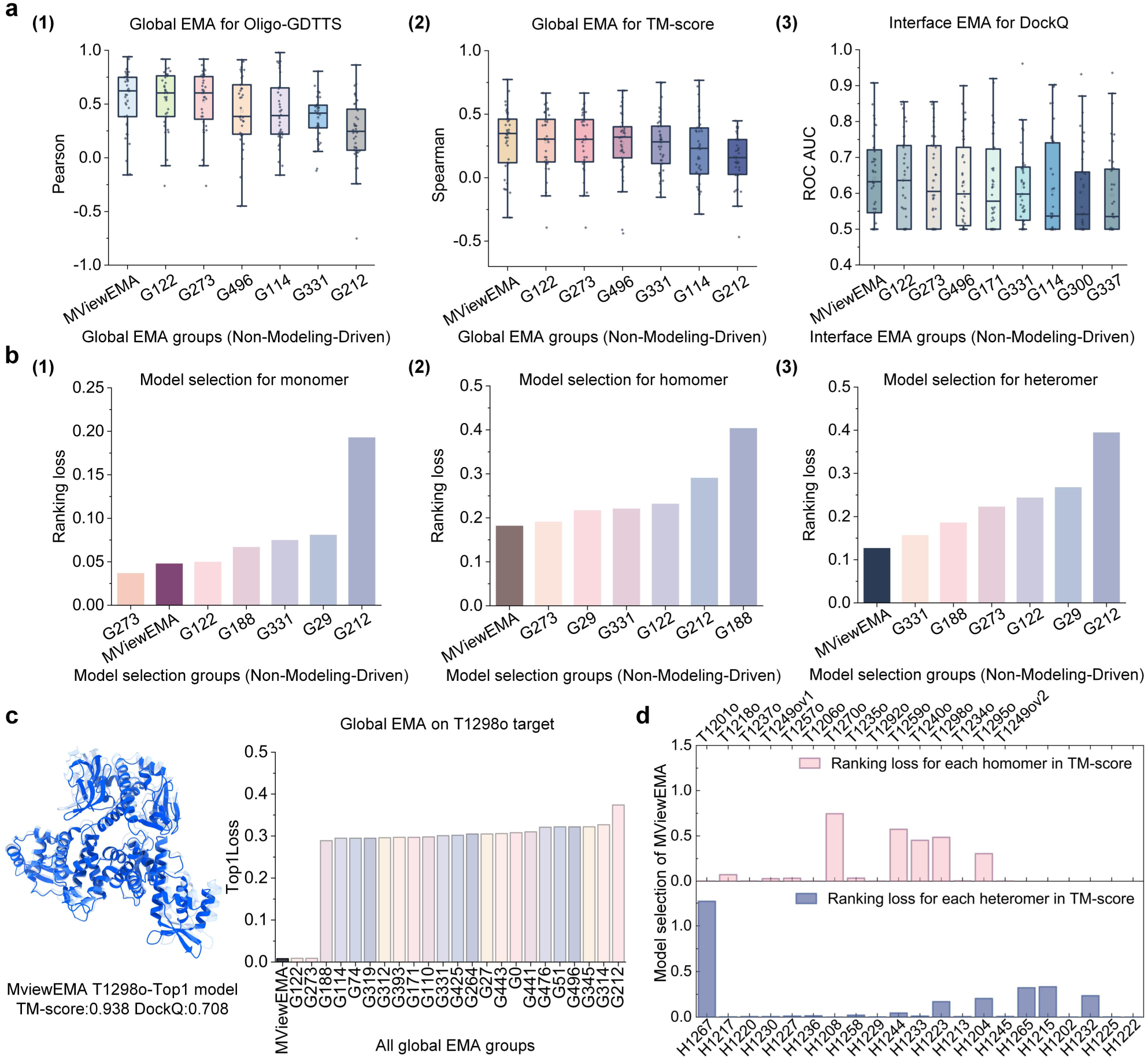
Performance comparison of MViewEMA and EMA methods using non-modeling-driven information in the CASP16 blind test. **a**, Comparison of global accuracy assessment performance between MViewEMA and other EMA methods based on (1) Pearson correlation of Oligo-GDTTS, (2) Spearman correlation of TM-score, and (3) ROC AUC of DockQ across CASP16 targets. **b**, TM-score ranking loss of MViewEMA and other EMA methods for monomer **(1)**, homomer **(2)**, and heteromer **(3)** model selection. **c**, Top1Loss of MViewEMA on homomer target T1298o. The native structure is shown in gray, while the selected structure is depicted in blue. **d**, TM-score ranking loss of MViewEMA for individual homomer (top panel) and heteromer (bottom panel) targets.

**Fig. 6.**
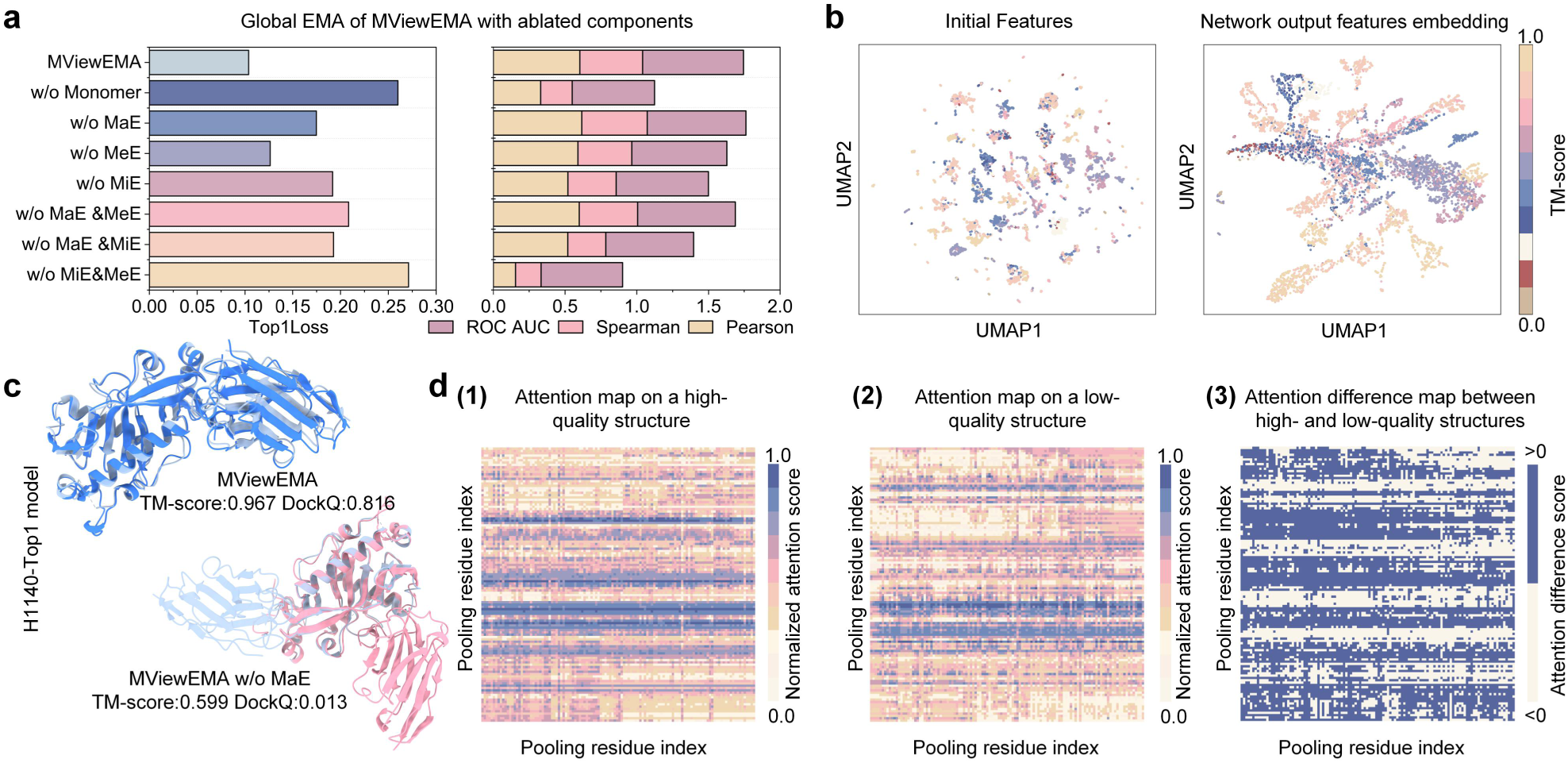
Component contribution analysis and network explainability of MViewEMA. **a**, Performance comparison of MViewEMA variants using different combinations of view features and training datasets on the benchmark dataset. **b**, UMAP visualizations of the initial features (left panel) and network-processed embeddings (right panel) from protein models. Each point is colored according to the corresponding model quality score (TM-score), with intervals from 0 to 1 in steps of 0.1. **c**, Structures of top-ranked models selected by MViewEMA and its w/o MaE variant on antibody-antigen target H1140. The native structure is shown in light blue, while the selected structures are depicted in distinct colors. **d**, Attention maps of high-quality **(1)** and low-quality **(2)** structures for target protein H1106, with attention scores normalized to [0, 1]. The heat map of the attention difference between high-quality and low-quality structures **(3)**, where blue regions indicate higher attention scores in high-quality structures, and white regions indicate higher attention scores in low-quality structures.

Interestingly, the global confidence scores predicted by MViewEMA also exhibited strong discriminative capability in assessing interface accuracy of protein models (**Fig. 5a (3)**). Specifically, MViewEMA achieved the best average ROC AUC (0.642) in distinguishing good and poor models under the DockQ metric. This suggests that despite the distinct focuses of various evaluation metrics, they may still provide complementary insights into model accuracy, potentially reflecting a correlation between inter-chain orientation and overall structural topology. This observation is consistent with correlation analysis between true DockQ and TM-score metrics, which yielded Pearson = 0.669 and Spearman = 0.798 (**Supplementary Fig. S9**). **Fig. 5c** further supports this observation by illustrating a blind test example of MViewEMA. On the homologous complex T1298o, MViewEMA achieved the highest ROC AUC (0.908) for interface accuracy assessment in the DockQ metric (**Supplementary Fig. S10**). Meanwhile, MViewEMA obtained the best Top1Loss (0.008) under the TM-score metric, significantly outperforming best single-model methods G312 (Top1Loss = 0.296) and G314 (Top1Loss = 0.327). Same result was also found in the Oligo-GDTTS metric (**Supplementary Table S9**). Based on the above results, we speculate that the MViewEMA may potentially identify the binding modes of complex protein targets.

For the MassiveFold^43^ model selection track, the models were generated by tuning the parameters of AlphaFold-Multimer, resulting in many highly similar conformations. To reduce computational overhead and improve efficiency of MViewEMA during the CASP16 experiment, we applied structural clustering to filter out redundant models prior to selection (see Methods for details), although this strategy may have affected overall model selection performance in the blind assessment. The blind test results, as evaluated by CASP assessors, analyzed the ranking loss of EMA methods between the predicted and observed top5 models for different types of protein targets, including monomer, homomer, and heteromer. It is worth noting that, since this study focuses on the analysis of global structural accuracy, all reported results are based on the TM-score metric (**Fig. 5b**). Analysis results based on other metrics can be found on the official website.

For monomer model selection, MViewEMA achieved a ranking loss of 0.048, which is behind G273 (0.037), as shown in **Fig. 5b (1)**. Among the 22 monomer targets, prediction results of MViewEMA were close to the best structures in 13 targets, with only 5 targets showing a ranking loss greater than 0.05 (**Supplementary Fig. S11**). This strong performance may be attributed to two main factors: (1) MViewEMA was trained using monomer structure datasets, enhancing its ability to evaluate monomer models; (2) existing structure prediction methods, particularly AlphaFold-based approaches, generated high-quality monomer models with minimal structural variability. For example, in monomer target T1276, the highest TM-score among protein models reached 0.968, while the lowest was still as high as 0.716. These results suggest that, from the perspective of EMA, the monomer protein structure prediction problem is largely resolved.

For homomer model selection, MViewEMA achieved the best ranking loss of 0.182 (**Fig. 5b (2)**), which was 3.79-fold worse than its monomer performance, potentially highlighting the substantial challenges of EMA that remain in complex model selection. On these targets, MViewEMA exhibited two distinct performance patterns, as shown at the top panel of **Fig. 5d**. It achieved near-optimal results on 10 targets, while showing suboptimal performance on 5 others, with a ranking loss exceeding 0.3. Particularly, for the challenging target T1270o, none of the EMA methods were able to identify high-quality models, with the best EMA method still showing a ranking loss of 0.586 (**Supplementary Fig. S12**). Further analysis of all 8,040 MassiveFold models for T1270o revealed that only 0.9% had a TM-score > 0.8, whereas 56.7% had a TM-score < 0.5. These findings suggest that, for challenging protein targets, existing EMA methods still struggle to identify reliable models from massive candidates in which only a small fraction are of high accuracy.

For heteromer model selection, MViewEMA outperforms other single-model methods with a ranking loss of 0.127, exhibiting better overall performance compared to its results on homomeric targets, as shown in **Fig. 5b (3)**. Notably, MViewEMA delivered near-optimal results on 15 out of 21 heteromer targets, indicating strong generalization ability across diverse heteromeric complexes, as shown at the bottom of **Fig. 5d**. However, its performance on target H1267 was limited, largely due to the challenging distribution of model quality for this target, with only 4.7% of models achieved a TM-score > 0.8, while 87% had TM-score < 0.5, a distribution pattern similar to that observed for target T1270o. For the challenging targets antibody–antigen complex, MViewEMA performed well on H1244, H1233, H1225, and H1222, achieving an average ranking loss of 0.015 and successfully selecting the best protein model for H1244. By contrast, it performed relatively poorly on H1215, H1232, H1204, and H1233 (average loss = 0.236), as shown in **Supplementary Table S10**. These results demonstrate MViewEMA has strong model assessment performance across diverse protein types, exhibiting well generalization in homomer and heteromer model selection.

### Ablation and explainability

To analyze the contribution of components in MViewEMA to global EMA performance, we conducted ablation studies using different view feature combinations and training datasets on the CASP15 benchmark dataset. Furthermore, we explored the explainability of deep neural networks in learning the mapping relationship between model features and structure accuracy, as illustrated in **Fig. 6**.

Specifically, the w/o Monomer variant (i.e., the MViewEMA variant excluding monomer model data during training) exhibited significantly lower performance across all evaluation metrics compared to the full version (**Fig. 6a**). This degradation underscores the critical role of monomer data for complex accuracy assessment, as it enables the network to learn fundamental folding principles and generalizable structural representations. This insight aligns with the training strategy adopted by AlphaFold-Multimer^42^, which leverages both monomer and complex data for effective structure modeling. For the w/o MaE variant, the MaE view features and their corresponding transform networks were removed, leaving only the MiE and MeE features and their networks to predict model quality. This resulted in a performance drop in Top1Loss to 0.175 (**Fig. 6a**). For some challenging protein targets, this variant selected low-quality structures instead of the optimal models, as observed in the antibody–antigen complex H1140 (**Fig. 6c** and **Supplementary Fig. S13**). These results suggest that incorporating the MaE view significantly enhances the network’s ability to identify high-quality structures by capturing long-range dependencies and global interaction patterns, which may be beneficial for characterizing the interaction relationships between cross-chain residues in complex models. To investigate the contribution of other view features, we conducted ablation studies on the MViewEMA by removing MiE, and MeE features, as well as their combinations. The results showed that all variants exhibit varying degrees of performance degradation across the analysis metrics. These findings highlight the importance of multi-view representation learning, which allows the network to capture structural patterns from diverse perspectives and thereby enhances the performance of global accuracy estimation.

To gain deeper insight into how MViewEMA evaluates protein structures, we visualized the transformation of feature embeddings in the network, as shown in **Fig. 6b**. Initial input features (i.e., prior to network processing) were extracted from all protein models and projected into two dimensions using UMAP^44^ tool. These initial features exhibit a sparse distribution with no apparent correlation to protein model quality (left panel of **Fig. 6b**). In contrast, the right panel of **Fig. 6b** illustrates that the output embeddings produced by the network become progressively clustered based on true quality scores, indicating that the network successfully learns the mapping between protein features and model quality. It is worth noting that, since the model quality labels are continuous values and the network’s prediction process is not error-free, the feature embeddings do not form completely independent clusters. Furthermore, for both high-quality (TM-score = 0.937) and low-quality (TM-score = 0.482) structures of H1106, the attention maps and corresponding attention scores obtained from the MViewEMA network enable us to analyze the network’s differential focus across various structural regions and how these differences impact the overall quality assessment, as shown in **Fig. 6d**. Compared to low-quality structures, high-quality structures exhibited heatmaps with more intense coloration and broader distribution, where attention scores have been normalized to the range [0,1]. To better distinguish the difference between high-quality and low-quality structures, **Fig. 6d (3)** shows their attention difference heat map, where the blue regions (i.e., the high-quality structure has higher attention scores) are significantly more extensive than the white regions (i.e., the low-quality structure has higher attention scores). This observation suggests that high-quality structures induce stronger and more widespread attention distributions associated with higher predicted scores, further demonstrating that the network effectively differentiates between high-quality and low-quality models.

## DISCUSSION

With the rapid advancement of deep learning technologies, the estimation of model accuracy has increasingly become more critical for modern protein structure prediction pipelines. Leveraging multi-view representation learning, we developed MViewEMA, a hierarchical deep learning framework for global accuracy assessment of protein models. Compared to existing EMA methods, the core advantage of MViewEMA lies in its introduction of three complementary views of residue-residue interactions: MiE, MeE, and MaE protein features, that are independent of modeling-driven information. It constructs corresponding sub-networks to systematically capture structural diversity and interactions across different scales. Extensive results demonstrate that MViewEMA achieves both high assessment accuracy and superior computational efficiency among single-model EMA methods, while also improving structure accuracy in mainstream deep learning-based prediction frameworks.

On the CASP15 benchmark, MViewEMA achieved the best global accuracy assessment performance among all single-model methods, with Z-score improvements of over at least 86.8% compared to our previous methods GraphGPSM and DeepUMQA3. In antibody – antigen complex evaluations, it also outperformed leading consensus approaches. Remarkably, MViewEMA was the only EMA method that successfully identified the best model for both challenging targets T1160o and T1161o. In the CASP16 blind test, MViewEMA achieved the best global Pearson and Spearman correlations among EMA methods without modeling-driven information. It demonstrated strong performance in interface-level accuracy assessment (ROC AUC) and maintained consistently high accuracy across monomers, homomers, and heteromers in the MassiveFold model selection, underscoring its robustness and practical value in structural biology research by leveraging multi-view representation learning.

For the current leading structure prediction method AlphaFold-Multimer, MViewEMA significantly improves accuracy of antibody-antigen models with its sampling strategies, resulting in an average TM-score increase of 10%. Moreover, MViewEMA achieved more accurate global accuracy estimates for protein complexes than the built-in self-assessment module of AlphaFold3. Importantly, although MViewEMA was not specifically trained on AlphaFold models, it exhibits strong generalizability and adaptability, making it a valuable complement to modern structure prediction pipelines. When integrated with the deep learning-based docking method DiffDock-PP, MViewEMA significantly improves structure accuracy, achieving 11.2% and 12.7% gains in TM-score and C-RMSD. This highlights not only the potential to enhance prediction performance but also the critical role of model selection strategies in obtaining high-quality structures. While the overall improvement was limited for the traditional template/energy-based docking method HDOCK, it achieves a notable 8.7% increase in TM-score on medium-quality models and can serve as an independent third-party tool to further validate the reliability of its model selections.

Recent CASP results suggest that protein structure prediction may be entering a model selection era, highlighting the growing importance of reliable EMA tools. As a next-generation protein EMA method, MViewEMA aims to offer the community a new perspective and evaluation framework to advance the EMA field and draw greater attention from the structural biology community. In future work, we plan to investigate interface-specific EMA methods for protein complexes and integrate high-performance modeling techniques to further improve model selection accuracy. Additionally, we will explore the scalability of MViewEMA to other biomolecules, such as DNA, RNA, and RNA–protein complexes.

## METHODS DETAILS

This section provides a detailed overview of the MViewEMA pipeline, including monomer and complex training datasets, multi-view features of protein models, and hierarchical network architecture. Additionally, we present details of the test datasets and describe the network training process.

### Training and test datasets

#### Training datasets

The training dataset of MViewEMA consists of protein monomer and complex model dataset. The monomer model dataset is derived from our previous work, the training dataset of GraphCPLMQA^24^ (with data before November 2021). Like the construction of the monomer dataset, the complex dataset is generated in three steps:

1. Complex target selection: A total of 11,838 protein dimers were selected from the PDB based on the following criteria: (a) minimum resolution <= 2.5 Å, (b) protein length 50-1000 residues, (c) sequence identity <= 40%, (d) and a release date before January 2022.
2. Model generation: For these dimers, decoy structures were generated using their native structures and ESMFold structures through xTrimoDock^45^, HDOCK, and native structure disturbance with chain orientation change.
3. Model sampling: The generated decoys were sampled based on their true DockQ scores [0, 1], selecting 10 to 25 models at intervals of 0.1, ultimately yielding a total of 1,443,856 complex protein structures.

#### Test datasets

The test dataset consists of four protein complex model datasets, including the CASP15 EMA benchmark dataset, the AlphaFold dataset, the Docking dataset, and the CASP16 EMA blind test dataset. These datasets (with data after May 2022) are strictly separated in time from the training set (with data before January 2022) to ensure the fairness and reliability of the experimental results. We adopt a temporal separation strategy to better simulate the real evaluation scenario of the predicted structure. Most of the test proteins are based on CASP targets, which is critical assessment of techniques for protein structure prediction.

*CASP15 EMA Benchmark Dataset* This dataset comprises predicted structures for 26 protein complex targets from CASP15 structure predicted groups, with approximately 350 models per target. These targets were chosen based on their suitability for AlphaFold-Multimer, AlphaFold3 and MViewEMA on a single NVIDIA A100 GPU, which is widely used for protein structure prediction and model accuracy assessment^46^ (**Supplementary Note S2**).

*AlphaFold Dataset* This dataset contains four types of AlphaFold-predicted models for CASP15 protein complex targets: (a) AFM Dataset-5: NBIS-AF2-multimer models provided by CASP, with five models per target. (b) AF3 Dataset-5: Structures generated by AlphaFold3 using default parameters, with five models per target. (c) AFM Dataset-MT: Structures generated by DeepSCFold^46^, based on AlphaFold-Multimer optimized MSAs, protein templates, and parameters with 6,015 models. (d) AF3 Dataset-Seeds: Structures generated by AlphaFold3 using 10 random seeds, with 1,300 models.

*Docking Dataset* This dataset includes two types of re-docking models based on native monomer structures for CASP15 protein dimers and PDB-filtered targets: (a) For protein dimer targets, the re-docking models were generated using the deep learning method DiffDock-PP and the template/energy-based method HDOCK, with 200 models per target, respectively. (b) For 116 protein targets filtered from the PDB (with release dates from 2024-01-01 to 2024-04-01, minimum resolution ≤ 2.5 Å, and protein length between 20 and 1,000), the re-docking models were generated using DiffDock-PP and HDOCK, with 200 models per target, respectively.

*CASP16 EMA Blind Test Dataset* The dataset consists of models from the structure prediction groups and models predicted by MassiveFold^43^: (a) Predicted structures for 38 protein complex targets submitted by CASP16 structure prediction groups, with approximately 350 models per target. These models are used to analyze the performance of the EMA methods in terms of global accuracy assessment (i.e., QMODE1).

(b) Predicted structures for protein 22 monomers and 36 complexes from MassiveFold, with 8,040 models per target (i.e., QMODE3). These models are employed to analyze the performance of the EMA methods in model selection. While there are 38 targets in QMODE1, model selection results of QMODE3 are unavailable for two of them in CASP official (https://predictioncenter.org/casp16/results.cgi).

### Multi-view features of protein

To combine multiple views to learn the relationship between protein structure and model accuracy, MViewEMA extracts geometric, sequence-based, and statistical energy features from the protein model and categorizes them into <u>mi</u>cro-<u>e</u>nvironment (MiE), <u>me</u>so-<u>e</u>nvironment (MeE) and <u>ma</u>cro-<u>e</u>nvironment (MaE) features based on residue – residue spatial relationships (**Supplementary Table S11**). This categorization establishes a progressive connection between local and global levels, providing three distinct views to characterize protein structures to enhance the accuracy and generalization of evaluation. It is worth noting that the core features of the method are its exclusive reliance on intrinsic protein features, excluding explicit evolutionary information (e.g., MSAs, templates or protein language embeddings). This design ensures that the prediction results are independent of protein structure modeling, offering a decoupled solution for protein model accuracy estimation.

#### Micro-environment (MiE) features

MViewEMA introduces amino acid properties, backbone geometric characteristics, secondary structure information (SS.), and the Rosetta one-body energy^47^ terms to represent the residue micro-environment:

1. Amino acid property features: For protein sequences, physicochemical (phys.-chem.) properties of the amino acids are extracted using Meiler^48^ and RDKit (http://www.rdkit.org), providing a quantitative characterization of their chemical and physical attributes. Additionally, one-hot encoding is used to represent different amino acid types.
2. Backbone geometric features: Torsion (i.e., Phi and Psi) angles, bond lengths (BB. lengths), and bond angles (BB. angles) are extracted from the protein backbone to characterize the protein micro-folding patterns and conformational states.
3. Secondary structure information: The secondary structure type (i.e., α-helix, β-sheet and coil) of each residue is identified to characterize the local geometric patterns and global fold organization of the protein, enabling the method to better capture the local conformational context of individual residues.
4. Rosetta one-body energy terms: Using the Rosetta energy function^47^, the sidechain-backbone interaction energy, Ramachandran conformation energy, backbone torsion angle energy, and Dunbrack side-chain conformation energy are calculated to assess the physical plausibility and stability of the protein models.

These MiE features primarily characterize the physicochemical properties, geometric attributes, and statistical potential information of individual residues, aiming to emphasize the distinct roles and structural contexts of each residue within a protein. This residue-specific micro-environmental information serves as a foundational layer for integrating MeE with MaE features.

#### Meso-environment (MeE) features

MViewEMA employs a voxel-based representation inspired by Ornate^49^ and DeepAccNet^29^ to capture the spatial geometric relationships between a residue and its surrounding atoms within the local environment. The voxelization process consists of the following steps:

1. Establish local environment: Each residue’s *C_α_* atom serves as the center to search for all heavy atoms within a *n* Å (n = 14) radius, thereby defining its local structural environment. To eliminate global rotation and translation effects in the environment, a local coordinate system is established based on the residue’s *C_α_*, *C_β_*, and *N* atoms, where all neighboring heavy atoms are transformed into this local environment ensuring equivariance of features.
2. Discretized coordinates: The *C_α_* atoms of the local environment were discretized by translation to the center of a *k* Å × *k* Å × *k* Å (*k* = 19.2) cubic grid that was uniformly divided into *m* × *m* × *m* (*m* = 24) voxels, where each neighboring atom is mapped onto a discrete voxel to grid the local spatial environment.
3. Trilinear interpolation: For neighboring atom (*x*, *y*, *z*), its position within the voxel grid is represented by the eight surrounding voxel vertices, with interpolation weights determined by its relative offset. This interpolation ensures smooth and continuous mapping of atomic positions in the 3D grid, reducing quantization artifacts and enhancing the spatial resolution of local structural features. To clearly describe the formula, the voxel size is set to 1:

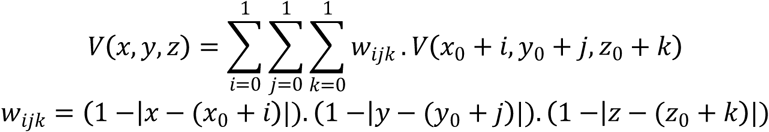

where *V*(*x*_0_ + *i*, *y*_0_ + *j*, *z*_0_ + *k*) is the voxel value at vertex (*x*_0_ + *i*, *y*_0_ + *j*, *z*_0_ + *k*).The voxel-based representation method provides a significant advantage in capturing local geometric variations of protein models, especially in protein complexes, where it facilitates the identification of geometric complementarity at interaction interfaces. Moreover, this structured 3-dim encoding serves as an input for deep learning networks such as 3D convolutional neural network, to efficiently learn spatial patterns within local environments, improving the EMA performance of protein complex. In addition, to enhance the perception of meso-scale residue – residue interactions beyond geometric level, MViewEMA integrates the BLOSUM62^38^ substitution matrix to represent the biochemical similarity and conserved substitutability of amino acids.

#### Macro-environment (MaE) features

MViewEMA utilizes distance and orientation maps between residues, residue frame alignment encoding, and Rosetta two-body energy terms^47^ to represent the macro-environment of each residue in relation to others:

1. Distance and orientation maps: For distance features, distance maps of different heavy atoms are extracted from the protein model, including *C_α_*, *C_β_*, and tip atoms, to capture the geometric distribution between each residue and others. For the orientation features, the *C_β_* atom of each residue was taken as the center, and other residues were sorted by *C_α_*-*C_β_* distance. Based on the order of neighboring residues, the dihedral angles between the residue and (i) all other residues, (ii) the side chains of other residues, and (iii) adjacent side chains are calculated, which helps to describe the spatial direction between the residues in the chains.
2. Residue frame alignment encoding: The geometric matrix of three translations and three rotations between each residue and all other residues is computed to represent the geometric constraints in the protein structure.
3. Rosetta two-body energy terms: Based on Rosetta energy function^47^, the interaction energy between each pair of residues is calculated and the presence hydrogen bonds are recorded to ensure that the interactions between residues conform to biological principles.

These MaE features establish a comprehensive representation of the global structural and energetic environment of among all residues to capture long-range dependencies and global interaction patterns, which is beneficial for characterizing the interaction relationship between cross-chain residues in complex models.

### Network architecture

MViewEMA adopts a hierarchical and heterogeneous neural architecture that facilitates information interaction across multi-scale structural levels of a protein model. It integrates graph attention networks (GATs)^34^, 3D convolutional neural networks (3D CNNs)^35^, and Transformers^36^, enabling comprehensive representation learning across MiE, MeE, MaE. GATs enable residue-level messages pass by combining MiE, MeE, and MaE features into a unified 1D representation. Meanwhile, 3D CNNs extract fine-grained geometric patterns from the local 3D environment (MeE), and Transformers integrate long-range dependencies between residues by processing MaE and MeE-2D features. These modules interact through shared representations and matrix operations, ultimately predicting a global confidence score for the protein model.

#### Graph attention networks

The GATs^34^ can assign attention weights to neighboring residues based on their geometric and physicochemical interactions. This adaptive weighting enables the network to focus on structurally critical regions, helping to identify the structural completeness of the protein. To further emphasize the network dynamic attention, the GATv2^50^ with dynamic weight is used in MViewEMA, which consists of the following two components (Fig. 1e):

1. Graph construction: The MiE-MeE-MaE representations are first pooled (pooling ratio = 0.5) to focus on part key residues that significantly impact overall structural accuracy. A protein graph G = (*V, E)* is then constructed by defining each residue’s *C_α_* atom as a node *v*_i_ ∈ *V* and identifying other *C_α_* atoms within a 15 Å radius to establish edges *e_ij_* ∈ *E*.
2. Cascaded GATv2 networks: Based on the protein graph *G*, the features ℎ*_i_* ∈ ℝ^𝑑^ serve as node inputs to a four-layer cascaded GATv2, where the layer employs 8 attention heads in parallel, generating microscopic residue embeddings and capturing diverse local structural information through iterative message passing between residues. The mathematical formula of the GATv2 layer can be described as follows:

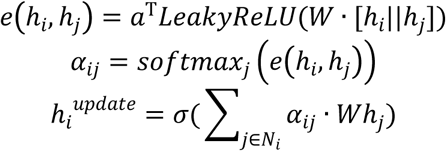

where *a* is a single-layer feedforward network used to calculate the attention score; *W* is shared linear transformation, parameterized by a weight matrix; *LeakyReLU* is employed as the nonlinear activation function; *𝑠oftmax_j_* means using the softmax function to normalize other residue node *j* attention coefficients; *σ* is a nonlinearity, such as a concatenation operation; The network consists of four layers with input channel dimensions of 77, 128, 128, and 128, and corresponding output dimensions of 16, 16, 16, and 8, respectively.

### 3D CNNs

The 3D CNNs^35^ offer unique advantages in processing the 3-dim data of protein structures, effectively capturing the geometric environment of local residues. The 3D CNNs in MViewEMA consist of the following three parts (Fig. 1c):

1. Representation downsampling: The voxelized MeE representation, with 167 channels corresponding to different atom types, are input into the 3D CNN layers, where the channel number of representation is sampled to 20, producing a latent residue-level structural descriptor.
2. Cascaded convolutions: For these descriptors, the first CNN layer (in_channels = 20, out_channels = 20, kernel_size = 3) extracts key local information; The second CNN layer (in_channels=20, out_channels = 30, kernel_size = 4) increases the number of channels (from 20 to 30), allowing for the extraction of spatial features across broader local regions; The third CNN layer (in_channels = 30, out_channels = 10, kernel_size = 4) reduces the channels (from 30 to 10) to decrease computational cost, resulting in a more condensed representation of the local structural context. This design balances feature extraction capability and computational efficiency.
3. Pooling and matching: The condensed representation undergoes a pooling operation (kernel_size = 4, stride = 4), which reduces spatial resolution while retaining key structural information. This is followed by horizontal striping, which symmetrically restructures the output into a MeE-2D representation, matching it with MaE features for efficient local-global integration in the subsequent transformer module.

#### Transformer Network

The transformer network^36^ with self-attention mechanism can capture the long-range dependencies in sequences or structures, thereby learning the global interactions between residues.

In MViewEMA, the transformer network module consists of the following three components (Fig. 1d):

1. Residues downsampling: The MaE representation fused with the MeE-2D and processed through a CNN layer (in_channels = 128*2, out_channels = 128) to generate inter-residue matrix that encodes pairwise spatial relationships, capturing both local interactions and the overall structural topology of the protein. Following this, the inter-residue matrix undergoes downsampling through a pooled convolutional network (in_channels = 128, out_channels = 128, kernel_size = 3, stride = 2, padding = 1), focusing on part residues that significantly impact the overall structure quality.
2. Biaxial attention: The downsampled matrix (*N, L, C*) is input into a multi-head attention (head = 4) mechanism network for both the *N* (pooling residue index) and *L* (pooling residue index) axes, each representing different residue indexing schemes, together with residue connectivity information. This design allows the network to focus on diverse interaction patterns across different space and residue dimensions, capturing complex topological dependencies and contextual relationships within protein structures from 4 distinct views. In addition, it can effectively prevent gradient vanishing or explosion.
3. Global accuracy prediction: The biaxial attention representation (i.e., MeE-MaE representation) and MiE-MeE-MaE-1D representation from the GAT network through dot product operation and average pooling are used to predict the global confidence score of the protein model.

### Train process

MViewEMA adopts a two-stage training strategy for regression prediction, utilizing the monomer and the complex model datasets (see training dataset) to train the network for predicting the global confidence score (i.e., TM-score) of protein models. Specifically, in the first stage (batch size = 1, 16-mixed), the network is trained for 50 epochs (each epoch with 7,552 steps) on the monomer model dataset to comprehensively learn the fundamental geometric features and structural patterns of protein models. In the second stage (batch size = 1, 16-mixed, each epoch with 11,838 steps), training transitions to the complex model dataset, with validation loss monitored to ensure convergence and stability. This progressive training strategy significantly enhances the network’s assessment performance, especially in capturing inter-chain interactions and complex topological dependencies in protein assemblies. To further improve the network’s adaptability to diverse data distributions, 95% of structural models are used as the training set and 5% as the validation set in each epoch, with decoy models randomly selected for training. The MViewEMA network employs the Log-cosh loss function^51^, defined as

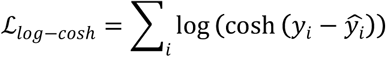

which combines the advantages of L1 and L2 losses by being smooth near zero and less sensitive to outliers. The method uses AdamW^52^ optimizer with parameters 𝑙 = 0.0001, β_1_ = 0.9, β_2_ = 0.999, *eps*= 1*e* − 08, *decoy* = 0.01. Training is performed on an NVIDIA A100-80G GPU, and the complete training procedure takes around three days.

## Structural clustering

To reduce computational overhead during CASP16, we randomly selected one structure from the 8,040 models generated by MassiveFold as a reference and aligned it against all remaining protein models using US-align^53^. The resulting structural similarity scores were binned at intervals of 0.001 to cluster the protein models. From each cluster, one representative structure was randomly selected to construct a candidate model pool for subsequent model selection.

## ACKNOWLEDGMENTS

This work was supported by the National Key R&D Program of China [2022ZD0115103], the National Nature Science Foundation of China [62173304], the “ Pioneer” and “ Leading Goose” R&D Program of Zhejiang [2025C01190], and the Zhejiang Province High-level Talent Special Support Program [2023R5248].

## AUTHOR CONTRIBUTIONS

Dong Liu (Formal Analysis, Methodology, Validation, Writing — original draft, Writing — review & editing), Xuanfeng Zhao (Methodology, Validation, Writing — review & editing), Tianyou Zhang (Formal Analysis, Validation), Lei Xie (Formal Analysis, Validation), Enjia Ye (Formal Analysis, Validation), Fang Liang (Formal Analysis, Validation), Haodong Wang (Formal Analysis, Validation), Guijun Zhang (Conceptualization, Formal Analysis, Methodology, Validation, Writing—review & editing).

## DECLARATION OF INTERESTS

The authors declare no competing interests.

## RESOURCE AVAILABILITY

### Lead contact

Further information and requests for resources and reagents should be directed to and will be fulfilled by the Lead Contact, Guijun Zhang.

### Data and code availability

The data supporting this study are available from the corresponding authors upon reasonable request. The release of inference code and model weights is currently under preparation.

## Supplementary Notes

**Supplementary Note S1** Sum Z-score formula in global accuracy estimation of CASP15

In global accuracy estimation of CASP15, all EMA methods are ranked based on group performance, using a combined assessor-based formula:

Sum Z-score=0.5 * Pearson (Oligo-GDTTS, Z-score) + 0.5 * Spearman (Oligo-GDTTS, Z-score) + AUC (Oligo-GDTTS, Z-score) – Top1Loss(Oligo-GDTTS, Z-score) + 0.5 * Pearson(TM-score, Z-score) + 0.5 * Spearman(TM-score, Z-score) + AUC(TM-score, Z-score) – Top1Loss(TM-score, Z-score).

**Supplementary Note S2** Protein targets of CASP15 EMA benchmark dataset

CASP15 protein targets were chosen based on their suitability for AlphaFold-Multimer, AlphaFold3 and MViewEMA on a single NVIDIA A100 GPU, which is widely used for large-scale protein structure prediction and model accuracy assessment: T1124o,T1121o,T1178o,T1153o,H1134,T1123o,T1179o,T1109o,T1110o,T1127o,T1113o,H1140,H1143,H11 42,H1141,H1144,T1187o,H1106,H1151,T1160o,T1161o,T1132o,T1173o,H1166,H1168,H1167.

## Supplementary Figures

**Supplementary Figure S1.**
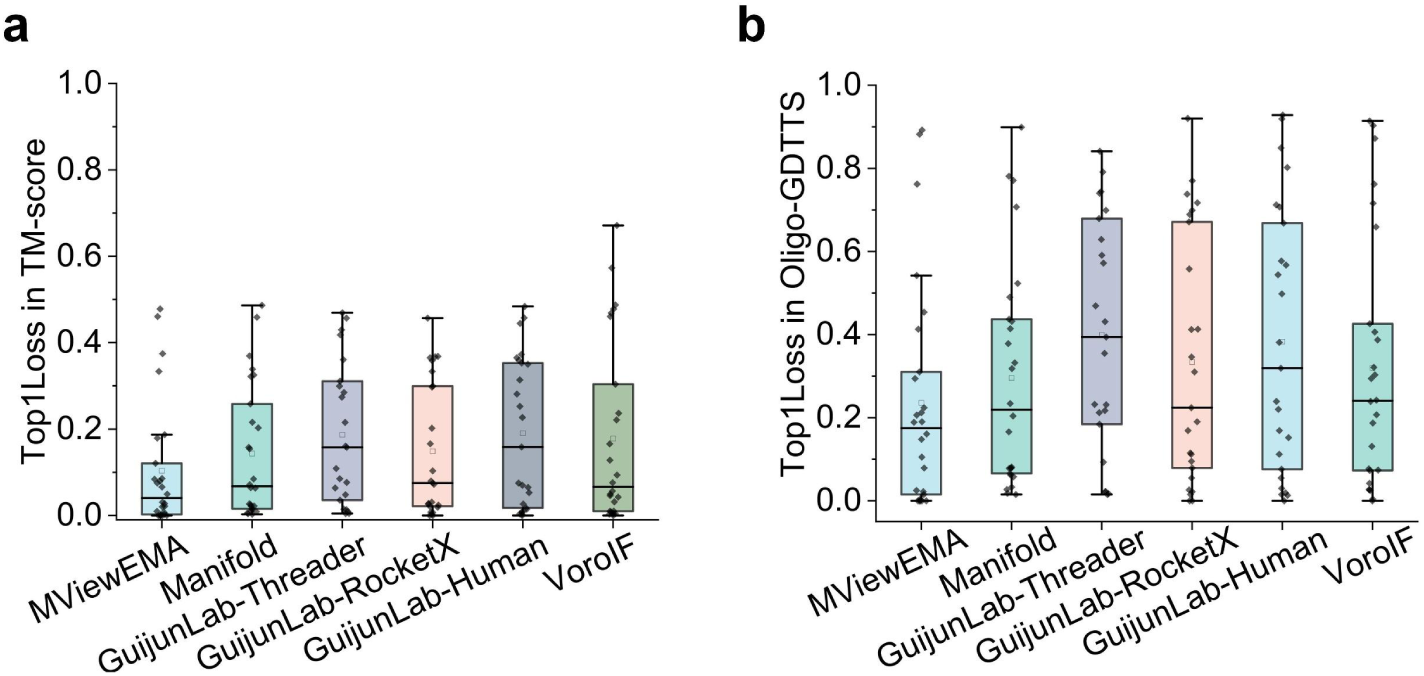
Top1Loss of MViewEMA and top 5 single-model methods on CASP15 models for the Oligo-GDTTS and TM-score metrics.

**Supplementary Figure S2.**
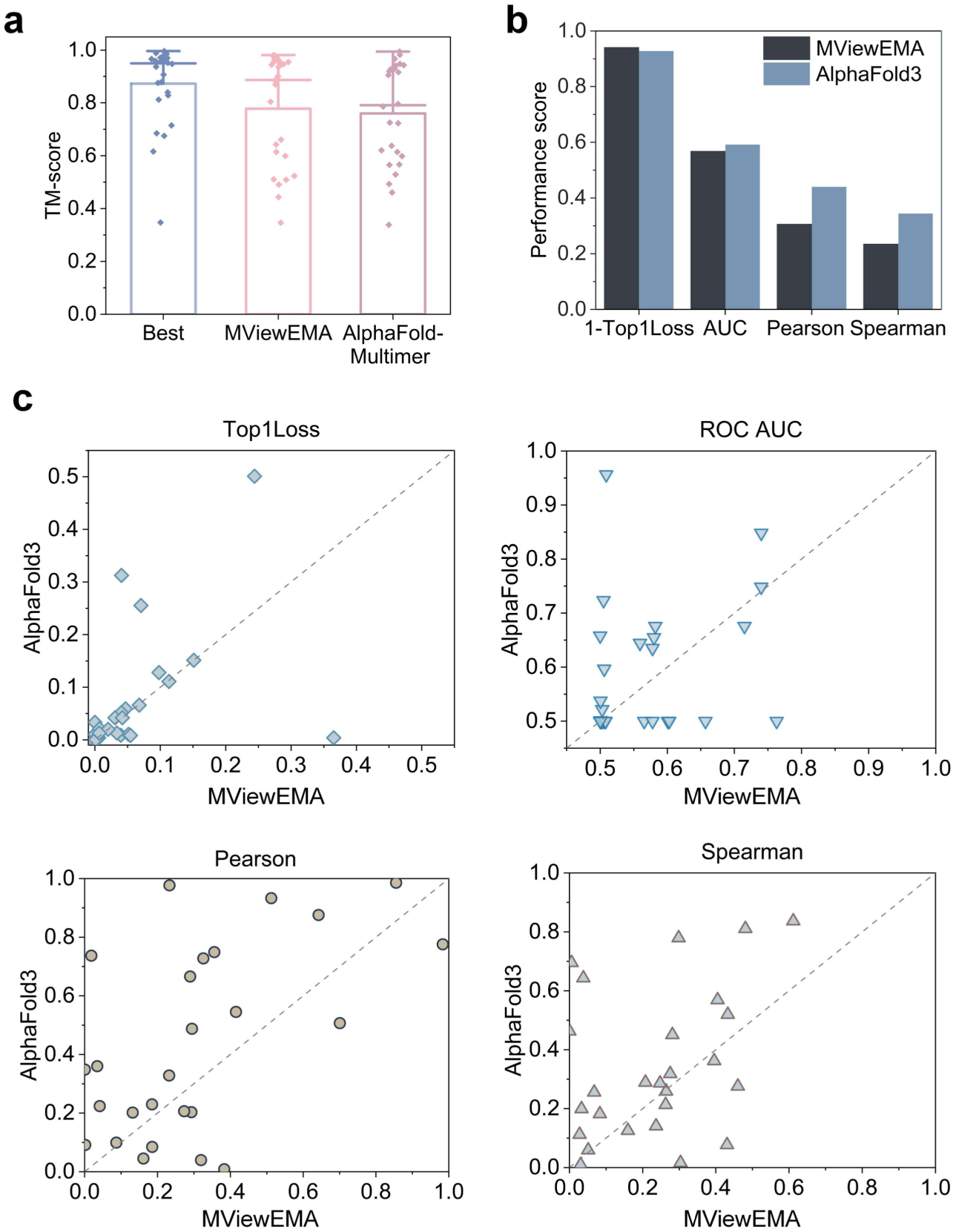
Performance comparison between MViewEMA and AlphaFold on five analytical metrics: TM-score (AFM), Top1Loss (AF3), ROC AUC (AF3), Pearson correlation (AF3), and Spearman correlation (AF3).

**Supplementary Figure S3.**
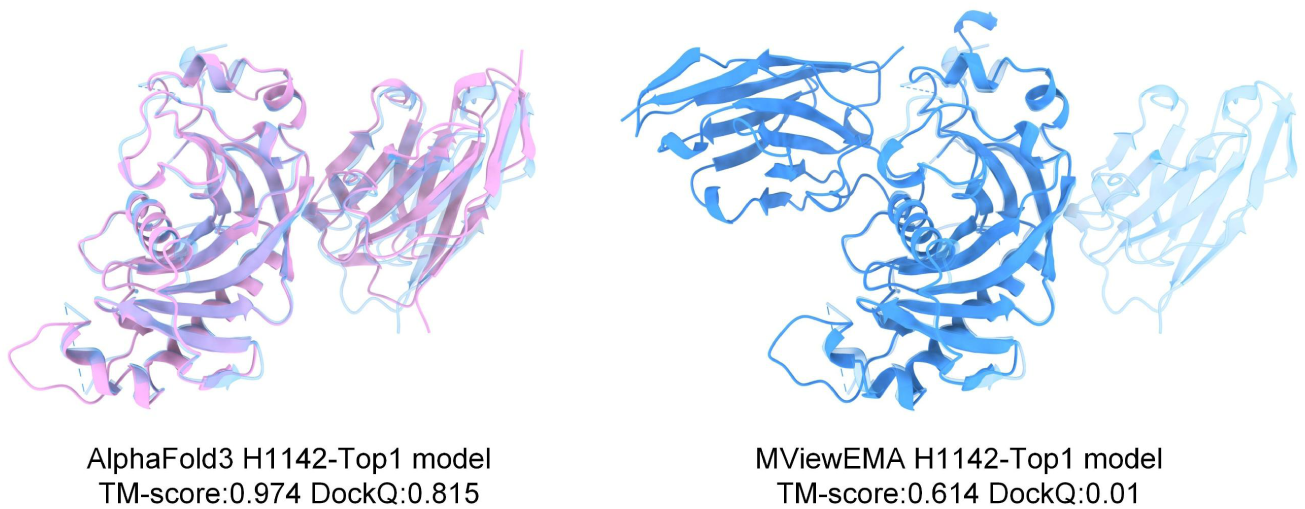
Top 1 model selected by MViewEMA and AlphaFold3 for antibody-antigen target H1142.

**Supplementary Figure S4.**
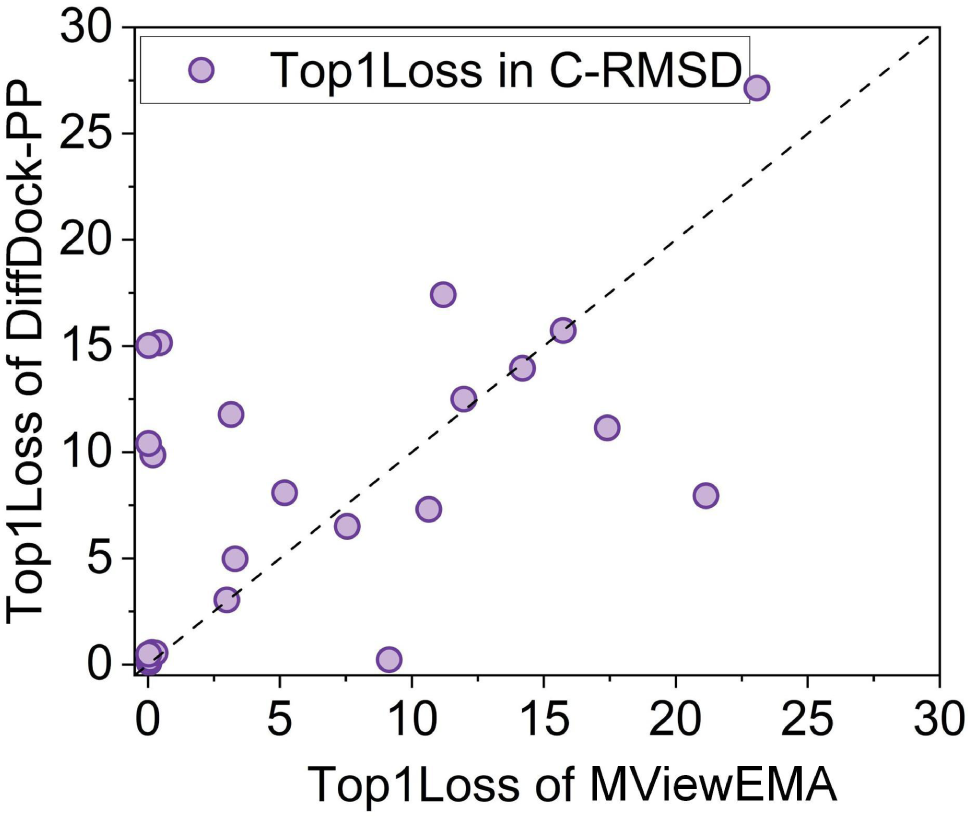
Top1Loss of the top1 models selected by MViewEMA and DiffDock-PP self-assessment on CASP15 dimers for C-RMSD metric.

**Supplementary Figure S5.**
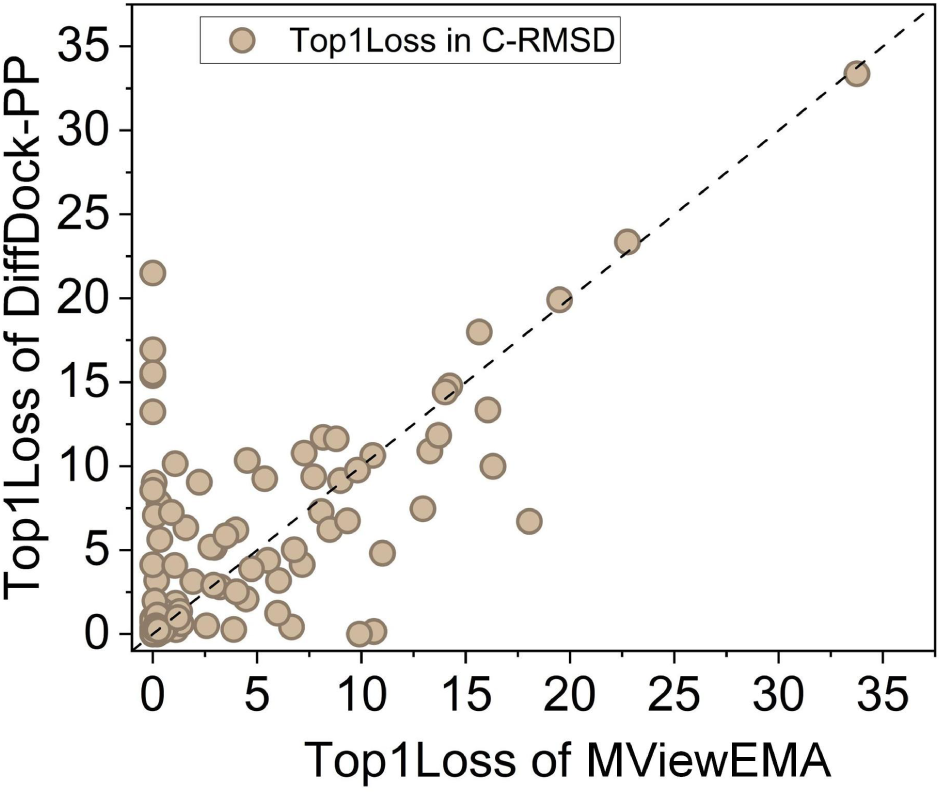
Top1Loss of the top1 models selected by MViewEMA and DiffDock-PP self-assessment on PDB-2024 targets for C-RMSD metric.

**Supplementary Figure S6.**
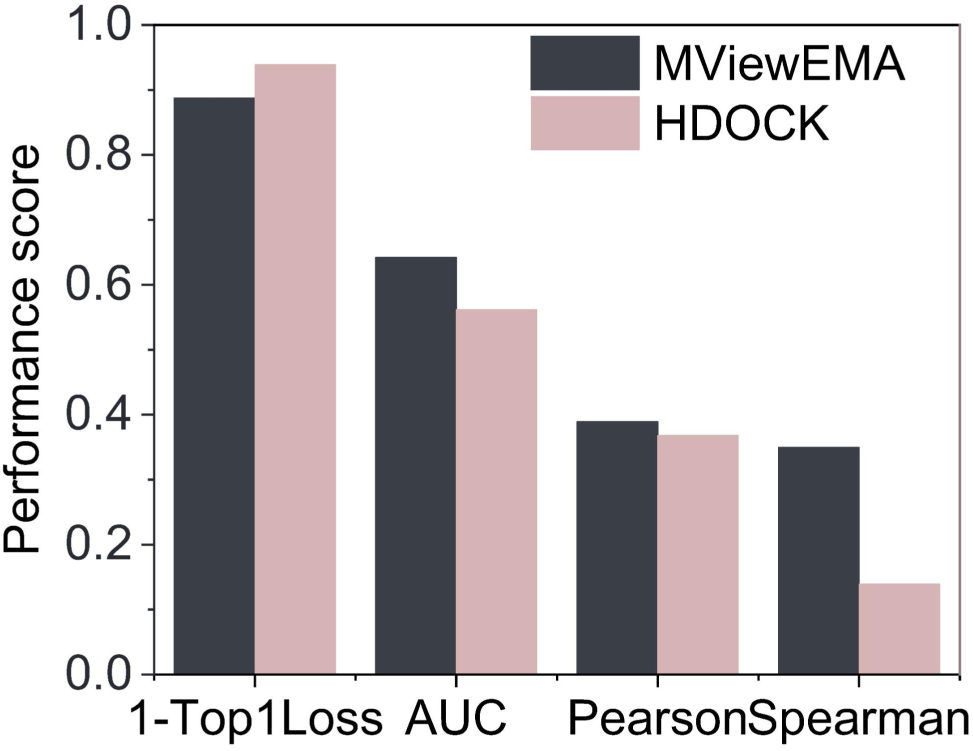
Performance comparison of MViewEMA and HDOCK self-assessment for CASP15 dimers.

**Supplementary Figure S7.**
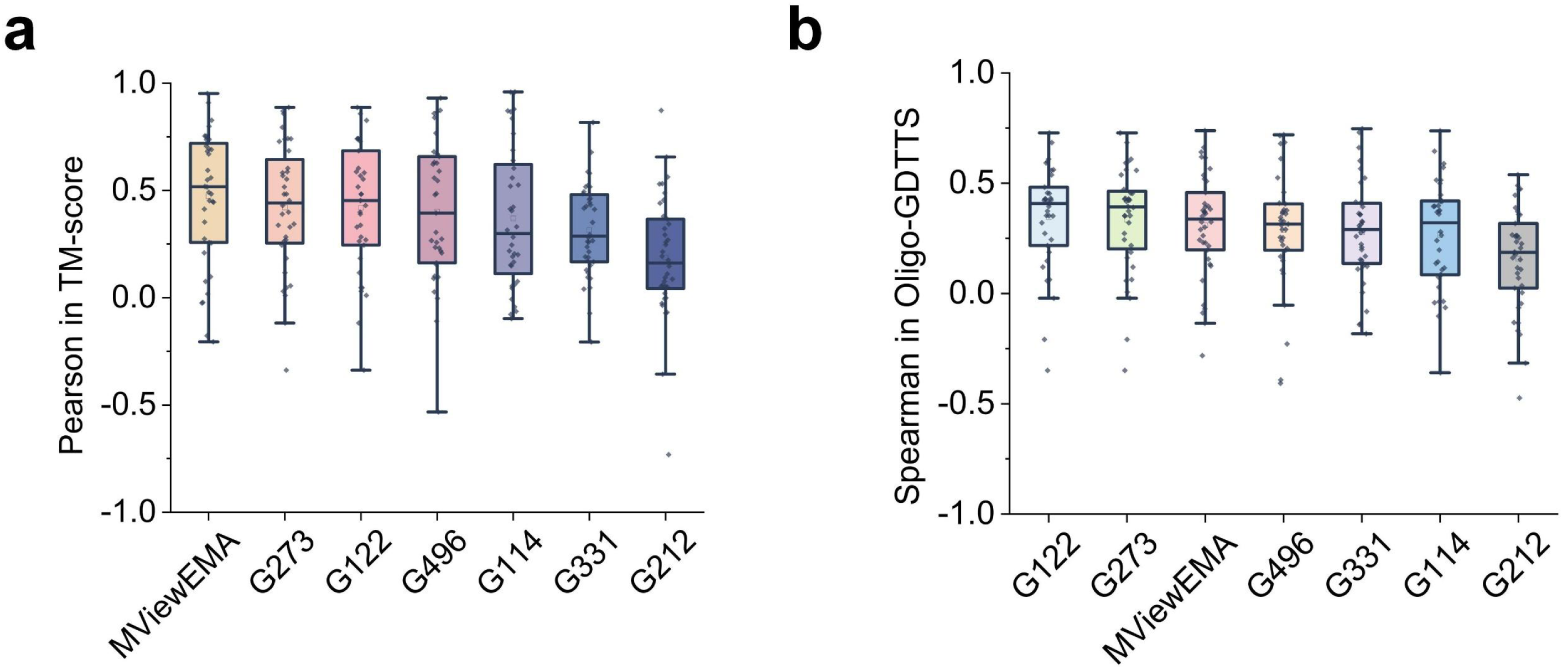
Correlation analysis of MViewEMA and all single-model methods that utilize non-modeling information in global accuracy assessment on the CASP16 blind dataset.

**Supplementary Figure S8.**
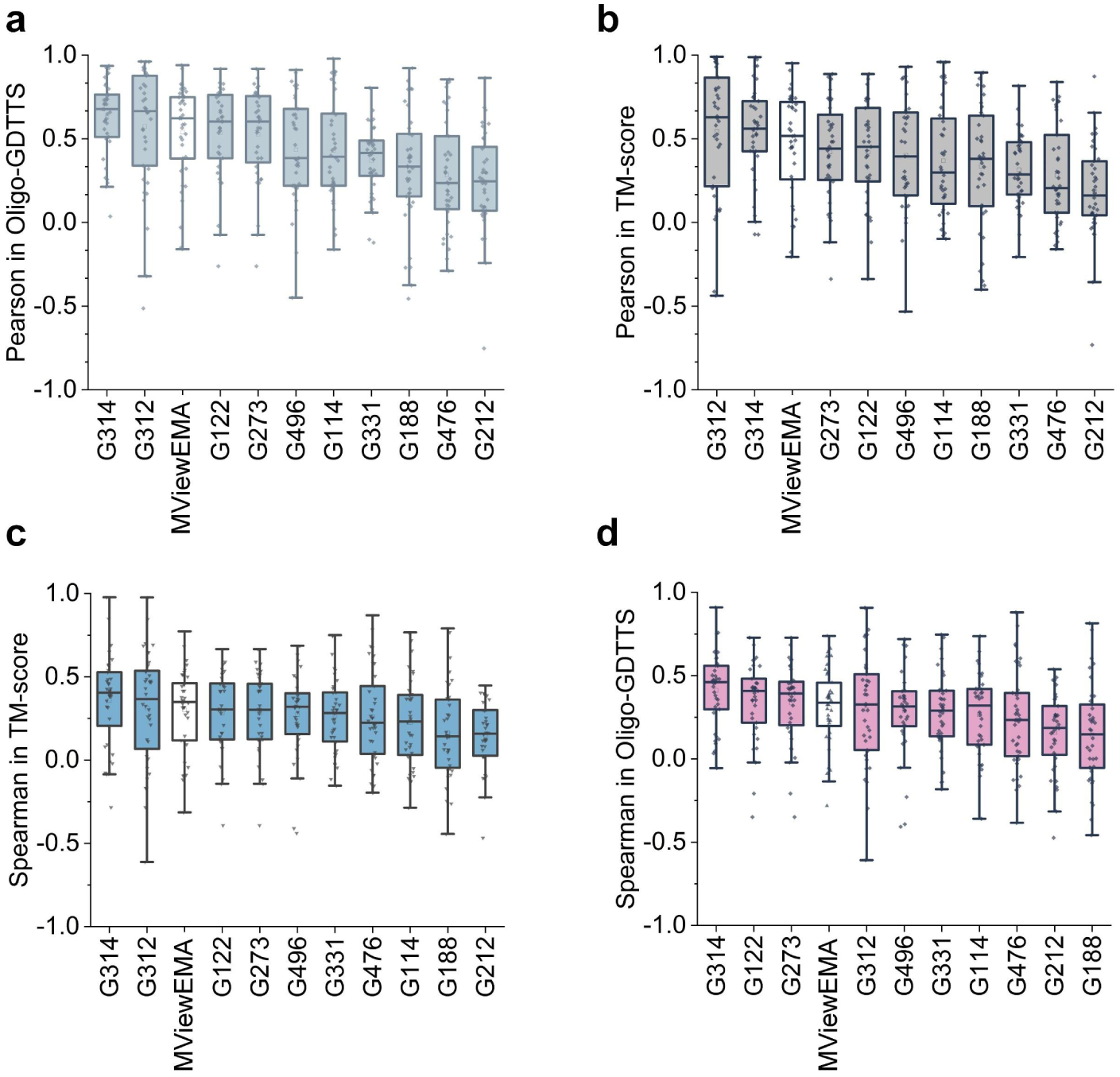
Correlation analysis of MViewEMA and all single-model methods in global accuracy assessment on the CASP16 blind dataset.

**Supplementary Figure S9.**
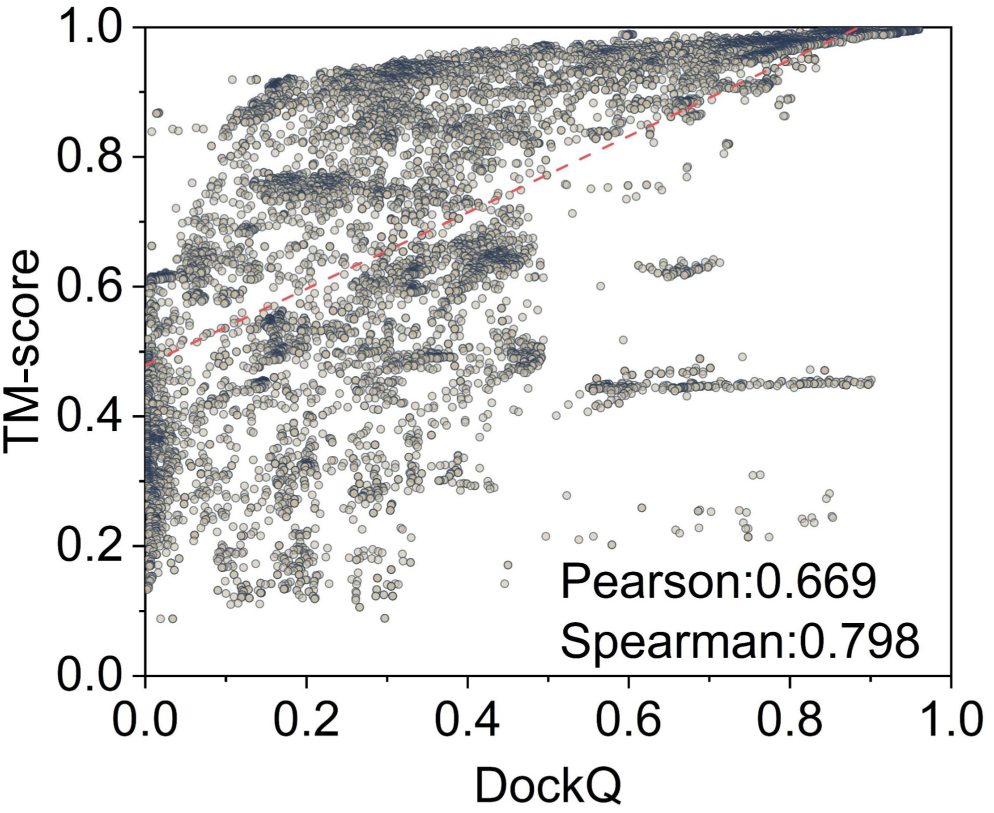
Ture quality distribution of interface quality DockQ and global quality TM-score on the CASP16 models.

**Supplementary Figure S10.**
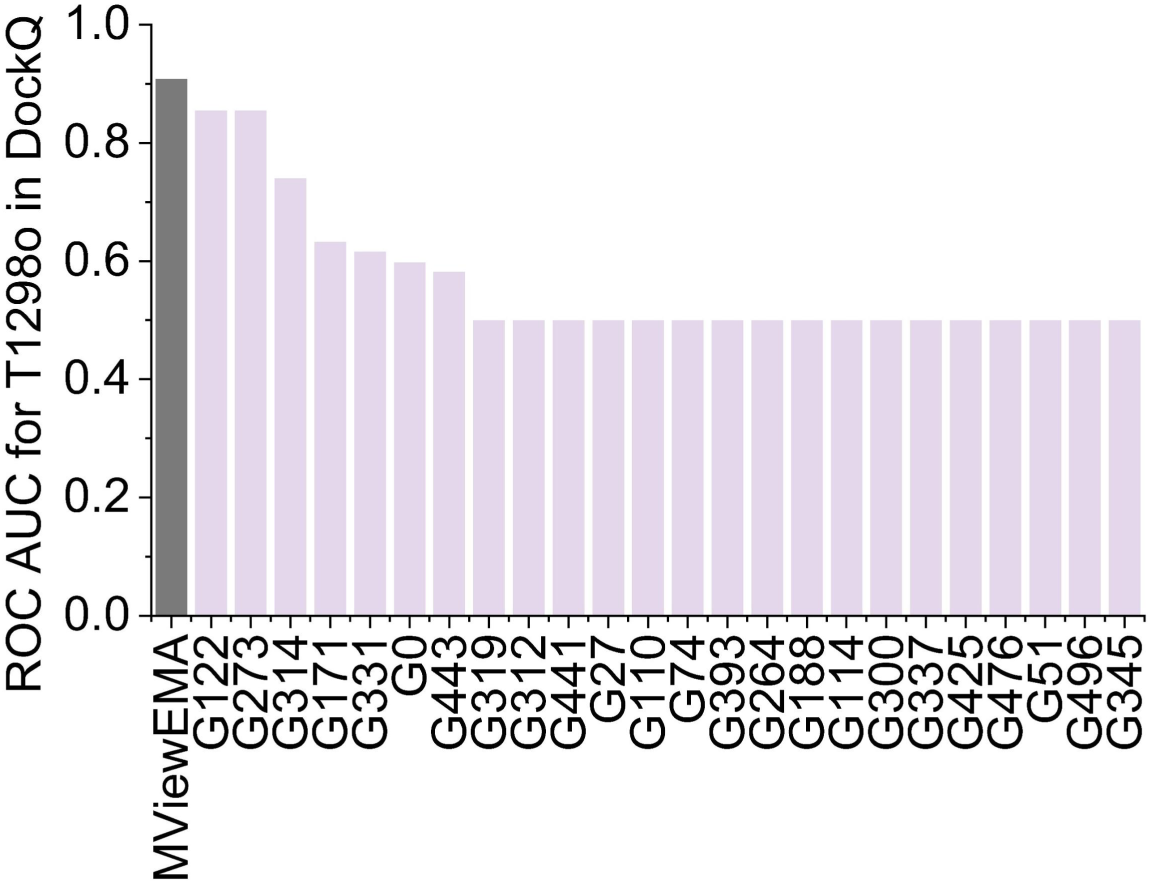
ROC AUC of DockQ for all EMA methods on target T1298o of CASP16.

**Supplementary Figure S11.**
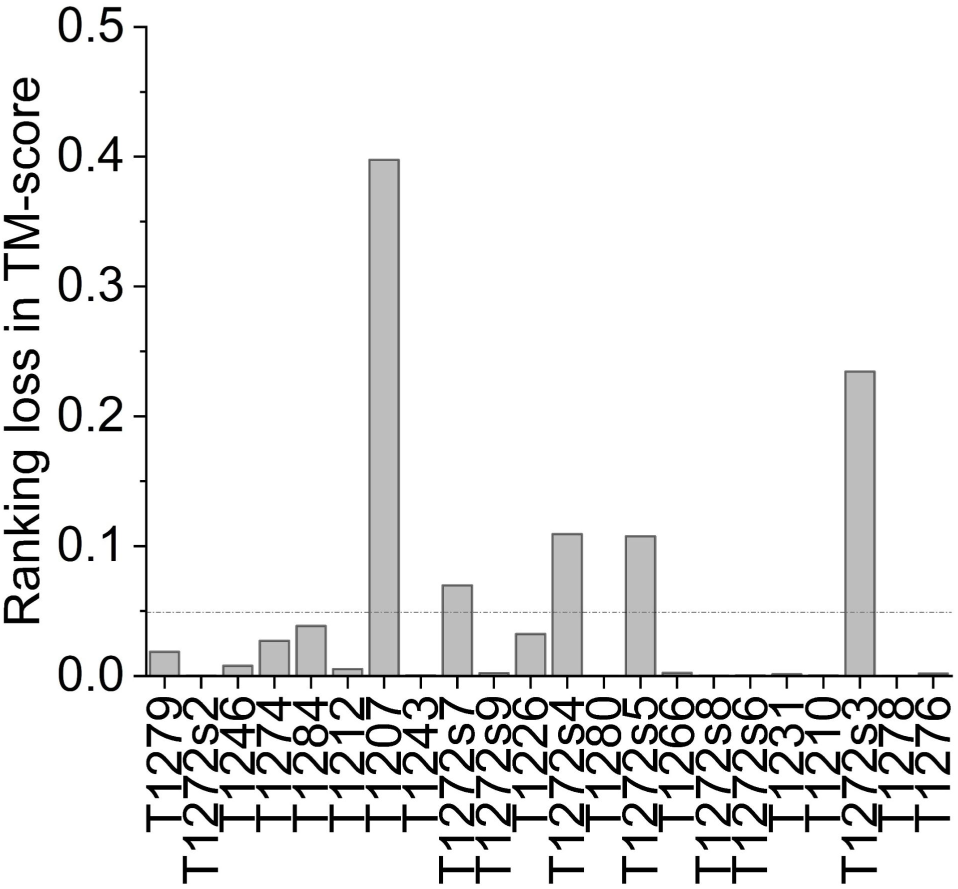
Ranking loss of MViewEMA in CASP16 monomer models.

**Supplementary Figure S12.**
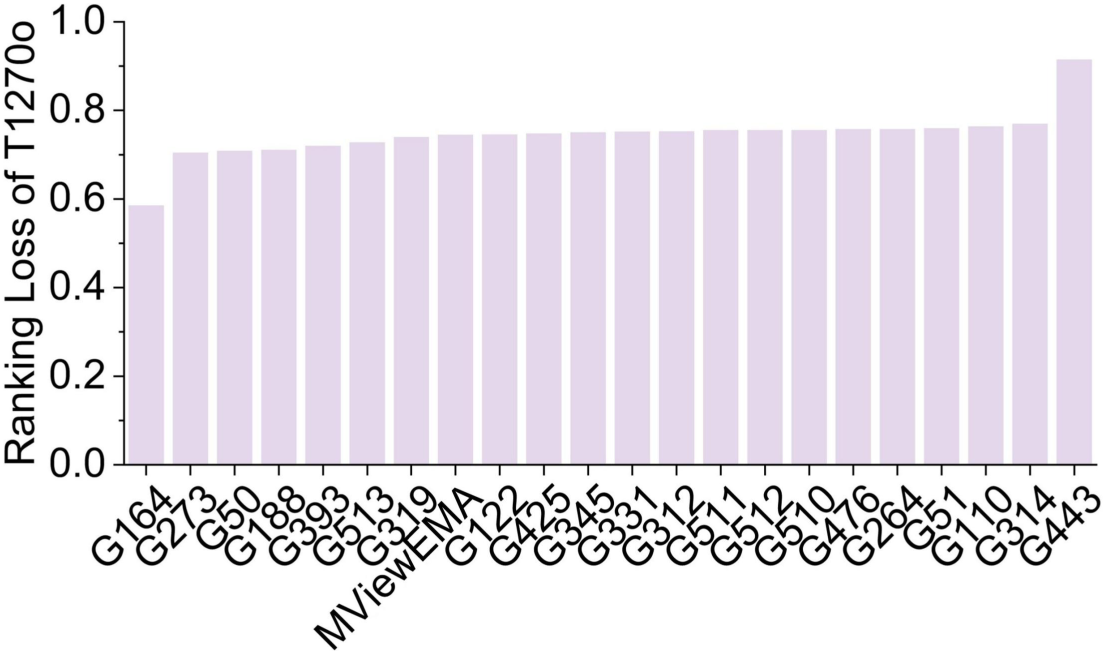
Ranking Loss of all EMA methods in homomer T1270o.

**Supplementary Figure S13.**
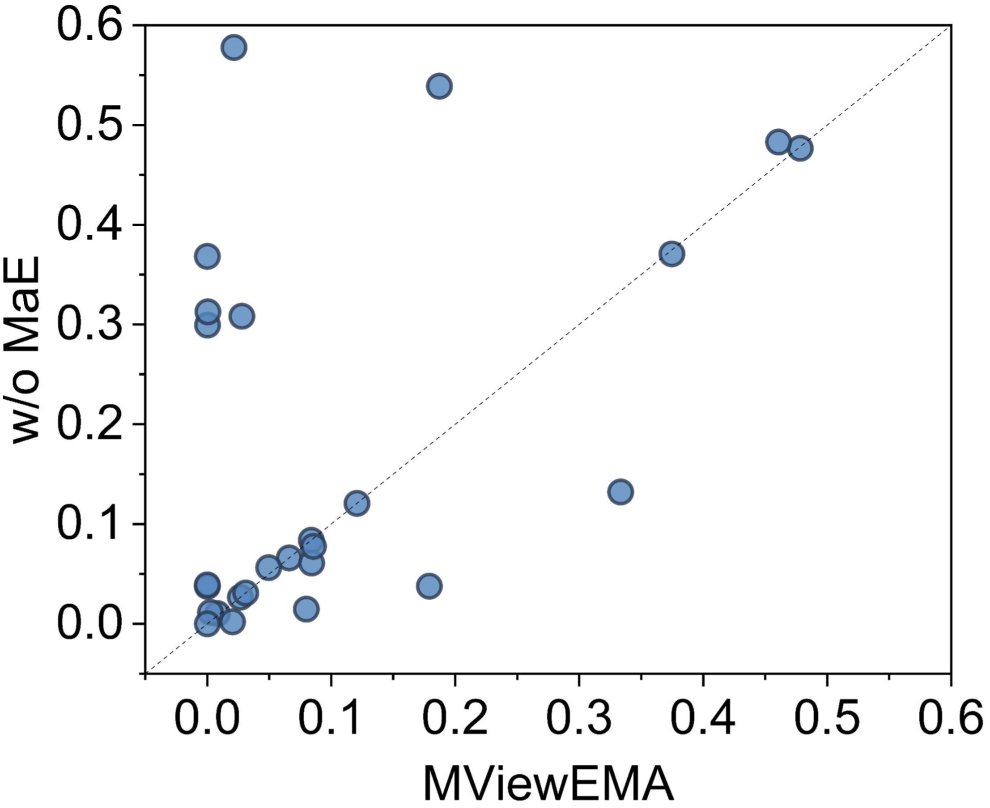
Top1Loss (TM-score) comparison between the full version MViewEMA and the w/o MaE variant on the CASP15 benchmark dataset

## Supplementary Tables

**Supplementary Table S1.** Performance of MViewEMA and all 16 single-model methods on CASP15 benchmark dataset in TM-score.

| Methods | Pearson | Spearman | ROC AUC | Top1loss |
| --- | --- | --- | --- | --- |
| MViewEMA | 0.603 | 0.438 | 0.704 | 0.104 |
| ChaePred | 0.424 | 0.312 | 0.644 | 0.282 |
| MUFold | 0.616 | 0.435 | 0.634 | 0.220 |
| MUFold2 | 0.556 | 0.328 | 0.603 | 0.241 |
| VoroIF | 0.500 | 0.353 | 0.637 | 0.178 |
| MULTICOM_egnn | 0.313 | 0.280 | 0.618 | 0.230 |
| VoroMQA-select-2020 | 0.403 | 0.387 | 0.659 | 0.196 |
| Manifold | 0.604 | 0.559 | 0.671 | 0.144 |
| MULTICOM_deep | 0.168 | 0.189 | 0.567 | 0.300 |
| GuijunLab-RocketX | 0.529 | 0.391 | 0.654 | 0.149 |
| GuijunLab-Human | 0.574 | 0.437 | 0.621 | 0.191 |
| GuijunLab-Assembly | 0.467 | 0.386 | 0.630 | 0.186 |
| MASS | 0.287 | 0.152 | 0.559 | 0.269 |
| LAW | 0.367 | 0.234 | 0.595 | 0.271 |
| APOLLO | 0.083 | 0.077 | 0.514 | 0.227 |
| GuijunLab-Threader | 0.657 | 0.538 | 0.669 | 0.187 |
| FoldEver | 0.413 | 0.262 | 0.597 | 0.224 |

**Supplementary Table S2.**
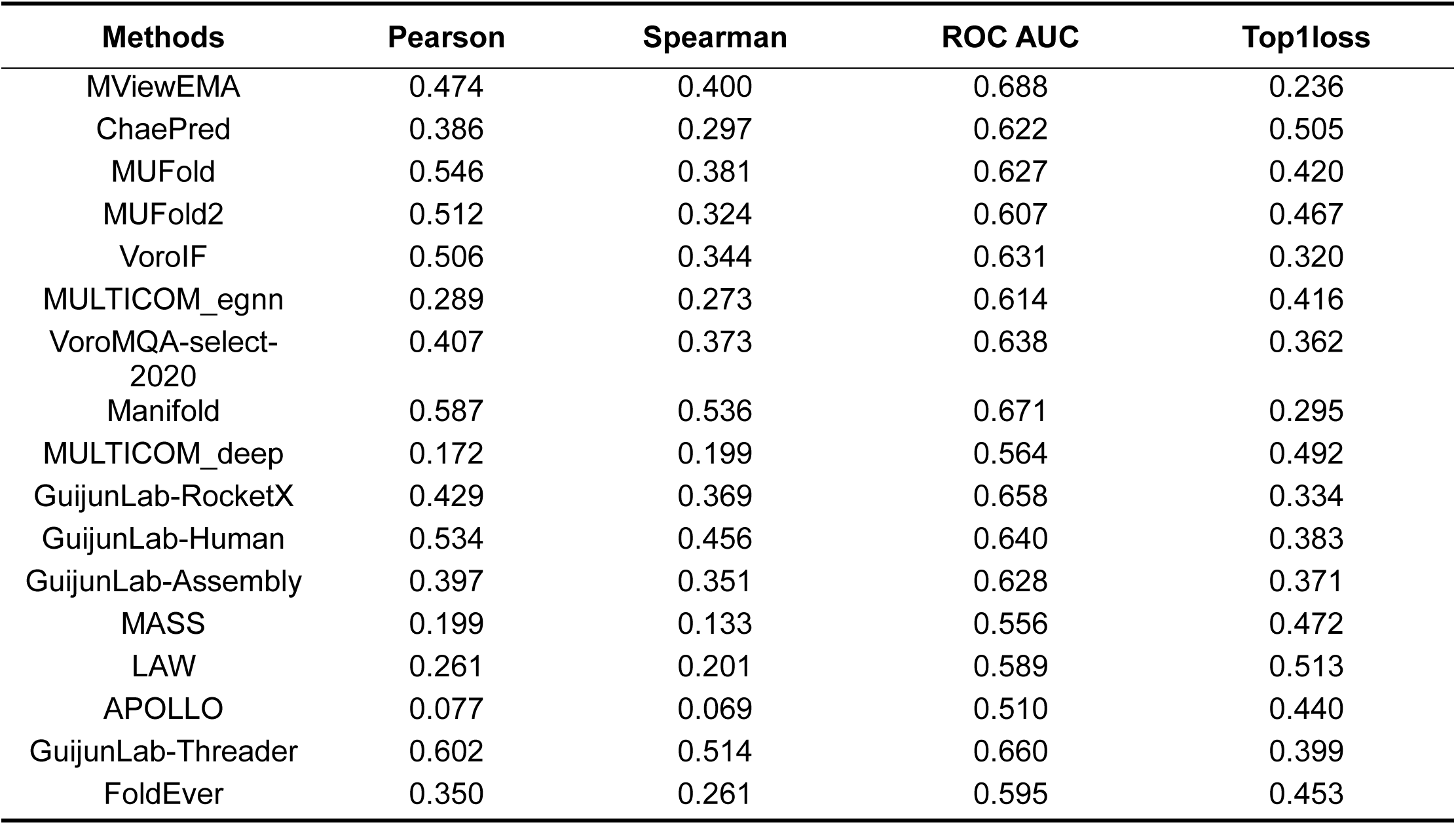
Performance of MViewEMA and all 16 single-model methods on CASP15 benchmark dataset in Oligo-GDTTS.

**Supplementary Table S3.** Comparison of model selection between MViewEMA and AlphaFold self-assessment in AFM dataset-5 and AF3 dataset-5 for TM-score.

| Dataset | Best | MViewEMA | AlphaFold |
| --- | --- | --- | --- |
| AFM Dataset-5 | 0.787 | 0.777 | 0.772 |
| AF3 Dataset-5 | 0.802 | 0.773 | 0.771 |

*Note: For each protein target in CASP15, AFM Dataset-5 and AF3 Dataset-5 benchmark datasets were* *constructed by generating five predicted structures using both AlphaFold-Multimer (from CASP15) and* *AlphaFold3(from AlphaFold3 server).*

**Supplementary Table S4.** Accuracy of the top1 models selected by MViewEMA and DiffDock-PP self-assessment on CASP15 dimers.

| Methods | TM-score | C-RMSD |
| --- | --- | --- |
| MViewEMA | 0.725 | 12.64 |
| DiffDock-PP | 0.652 | 14.48 |
| Best | 0.789 | 5.78 |

**Supplementary Table S5.**
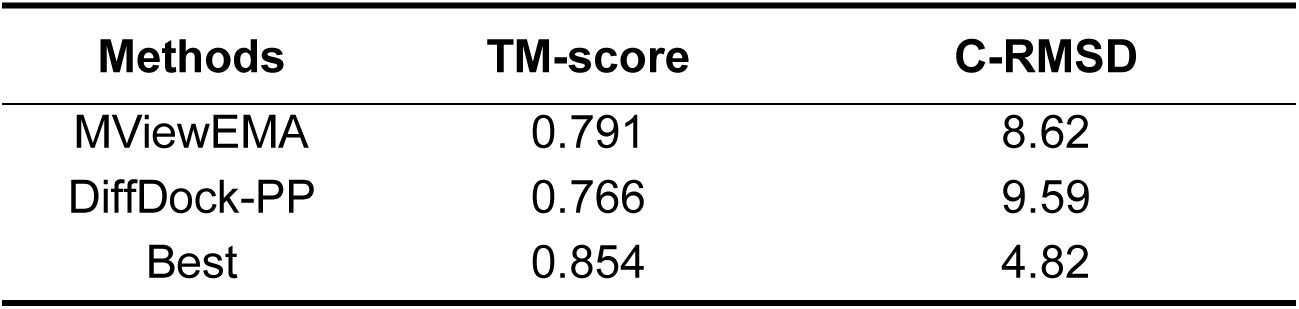
Accuracy of the top1 models selected by MViewEMA and DiffDock-PP self-assessment on PDB-2024 targets.

**Supplementary Table S6.**
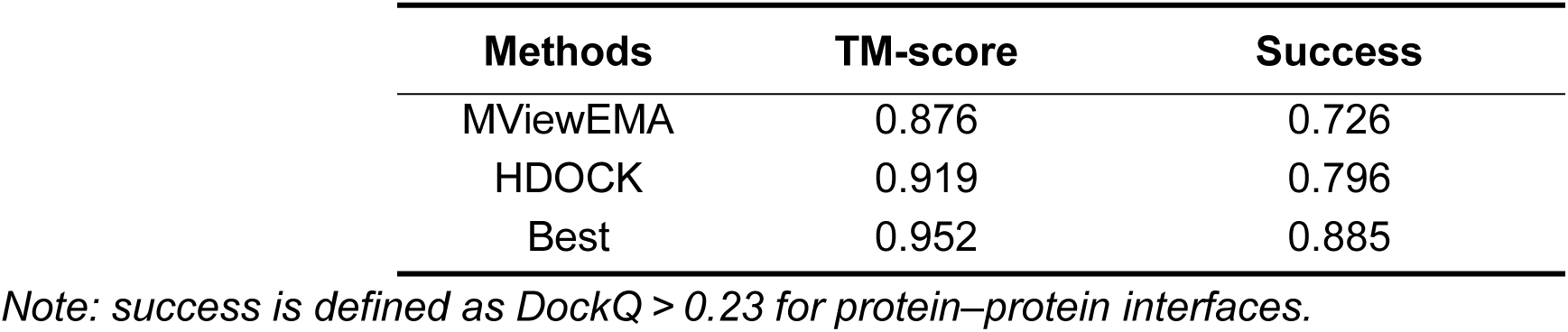
Accuracy of the top1 models selected by MViewEMA and HDOCK self-assessment on PDB-2024 targets.

| Methods | TM-score | Success |
| --- | --- | --- |
| MViewEMA | 0.876 | 0.726 |
| HDock | 0.919 | 0.796 |
| Best | 0.952 | 0.885 |

*Note: success is defined as DockQ > 0.23 for protein–protein interfaces.*

**Supplementary Table S7.** Accuracy of the top1 models selected by MViewEMA and HDOCK self-assessment on 12 medium-quality targets for TM-score metric.

| Targets | HDock | MViewEMA |
| --- | --- | --- |
| 8YE5 | 0.526 | 0.527 |
| 8XLD | 0.156 | 0.649 |
| 8UAC | 0.512 | 0.514 |
| 8QF3 | 0.556 | 0.511 |
| 8PZH | 0.513 | 0.513 |
| 8PQP | 0.528 | 0.527 |
| 8K89 | 0.527 | 0.548 |
| 8IMF | 0.521 | 0.529 |
| 8COP | 0.514 | 0.572 |
| 8CM8 | 0.506 | 0.499 |
| 7YP0 | 0.542 | 0.542 |
| 7YMF | 0.636 | 0.636 |

**Supplementary Table S8.** Comparison of MViewEMA with the best single-methods for CASP16 antibody-antigen targets in Pearson correlation (TM-score metric) of global accuracy estimation.

| Targets | MViewEMA | G314 | G312 |
| --- | --- | --- | --- |
| H1204 | -0.022 | -0.072 | 0.216 |
| H1215 | 0.274 | 0.253 | 0.158 |
| H1222 | 0.683 | 0.680 | 0.580 |
| H1223 | 0.353 | 0.319 | -0.413 |
| H1225 | 0.670 | 0.596 | 0.497 |
| H1232 | 0.491 | 0.479 | 0.526 |
| H1233 | 0.448 | 0.554 | 0.069 |
| H1244 | 0.738 | 0.581 | 0.623 |

**Supplementary Table S9.** Comparison of Top1Loss between MViewEMA and all EMA methods on T1298o models of CASP16 for Oligo-GDTTS.

| Methods | Top1Loss | Methods | Top1Loss | Methods | Top1Loss |
| --- | --- | --- | --- | --- | --- |
| MViewEMA | 0.032 | G319 | 0.21 | G476 | 0.255 |
| G122 | 0.037 | G171 | 0.212 | G27 | 0.26 |
| G273 | 0.037 | G110 | 0.214 | G51 | 0.266 |
| G314 | 0.091 | G425 | 0.218 | G496 | 0.266 |
| G114 | 0.2 | G0 | 0.221 | G345 | 0.266 |
| G331 | 0.205 | G188 | 0.224 | G212 | 0.394 |
| G393 | 0.206 | G264 | 0.227 |  |  |
| G312 | 0.208 | G443 | 0.229 |  |  |
| G74 | 0.21 | G441 | 0.252 |  |  |

**Supplementary Table S10.**
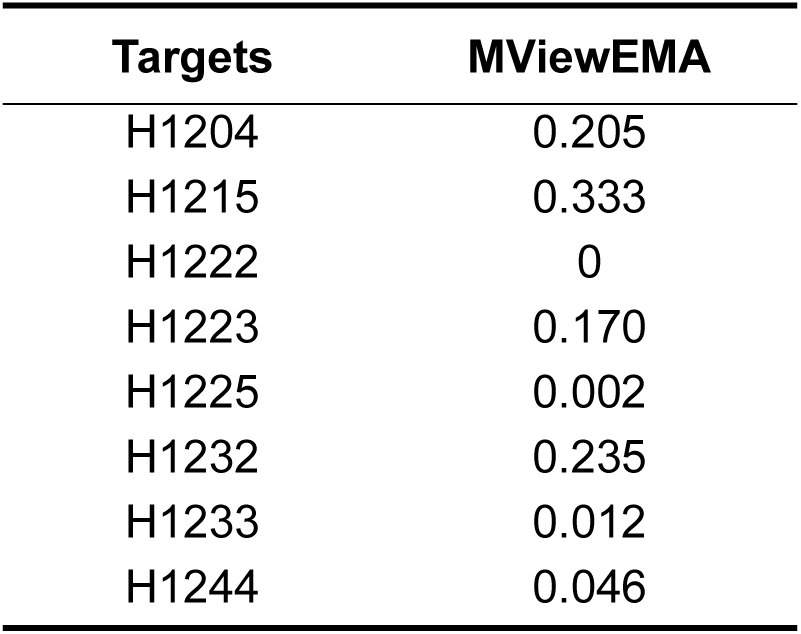
Ranking loss of MViewEMA for MassiveFold models of CASP16 antibody-antigen in TM-score.

| Targets | MViewEMA |
| --- | --- |
| H1204 | 0.205 |
| H1215 | 0.333 |
| H1222 | 0 |
| H1223 | 0.170 |
| H1225 | 0.002 |
| H1232 | 0.235 |
| H1233 | 0.012 |
| H1244 | 0.046 |

**Supplementary Table S11.**
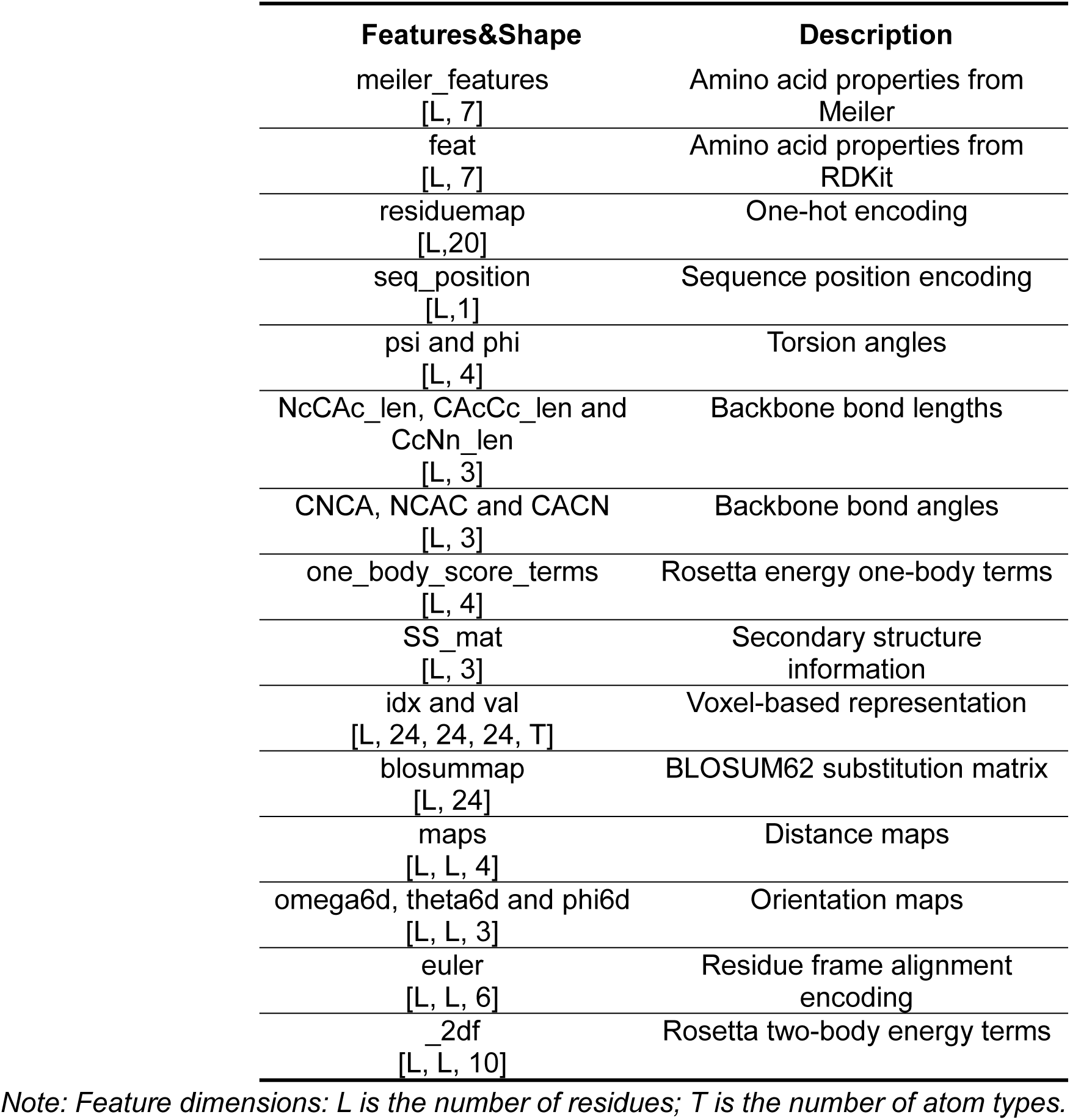
Input features of MViewEMA.

| Features&Shape | Description |
| --- | --- |
| meiler_features<br>[L, 7] | Amino acid properties from Meiler |
| feat<br>[L, 7] | Amino acid properties from RDKit |
| residuemap<br>[L,20] | One-hot encoding |
| seq_position<br>[L,1] | Sequence position encoding |
| psi and phi<br>[L, 4] | Torsion angles |
| NcCAc_len, CAcCc_len and CcNn_len<br>[L, 3] | Backbone bond lengths |
| CNCA, NCAC and CACN<br>[L, 3] | Backbone bond angles |
| one_body_score_terms<br>[L, 4] | Rosetta energy one-body terms |
| SS_mat<br>[L, 3] | Secondary structure information |
| idx and val<br>[L, 24, 24, 24, T] | Voxel-based representation |
| blosummap<br>[L, 24] | BLOSUM62 substitution matrix |
| maps<br>[L, L, 4] | Distance maps |
| omega6d, theta6d and phi6d<br>[L, L, 3] | Orientation maps |
| euler<br>[L, L, 6] | Residue frame alignment encoding |
| _2df<br>[L, L, 10] | Rosetta two-body energy terms |

*Note: Feature dimensions: L is the number of residues; T is the number of atom types.*

## Notes

### Competing Interest Statement

The authors have declared no competing interest.

